# Deterministic and stochastic processes generating alternative states of microbiomes

**DOI:** 10.1101/2023.04.03.535343

**Authors:** Ibuki Hayashi, Hiroaki Fujita, Hirokazu Toju

**Affiliations:** Center for Ecological Research, Kyoto University, Otsu, Shiga 520-2133, Japan

**Keywords:** biodiversity, community assembly, microbiome dynamics, ecological drift, alternative stable states, multiple stability, transient dynamics, dysbiosis

## Abstract

The structure of microbial communities is often classified into discrete or semi-discrete compositions of taxa as represented by “enterotypes” of human gut microbiomes. Elucidating mechanisms that generate such “alternative states” of microbiome compositions has been one of the major challenges in ecology and microbiology. In a time-series analysis of experimental microbiomes, we here show that both deterministic and stochastic ecological processes drive divergence of alternative microbiome states. We introduced species-rich soil-derived microbiomes into eight types of culture media with 48 replicates, monitoring shifts in community compositions at six time points (8 media × 48 replicates × 6 time points = 2,304 community samples). We then confirmed that microbial community structure diverged into a few discrete states in each of the eight medium conditions as predicted in the presence of both deterministic and stochastic community processes. In other words, microbiome structure was differentiated into a small number of reproducible compositions under the same environment. This fact indicates not only the presence of selective forces leading to specific equilibria of community-scale resource use but also the influence of demographic drift (fluctuations) across basins of such equilibria. A reference-genome analysis further suggested that the observed alternative states differed in ecosystem-level functions. These findings will help us examine how microbiome structure and functions can be controlled by changing the “stability landscapes” of ecological community compositions.

## INTRODUCTION

Understanding the mechanisms by which microbial community structure is organized is a major challenge in ecology and microbiology [1–4]. In a general framework of community dynamics, selection, diversification (speciation), dispersal, and drift have been considered as fundamental components of community processes [5, 6] (Fig. 1). Among the four component processes [5, 6], selection and drift can be regarded, respectively, as purely deterministic and stochastic components [7, 8], while dispersal and diversification are considered to include both deterministic and stochastic components [8]. Elucidating how those deterministic and stochastic processes collectively organize ecological community assembly is the key to predict and manage microbiome dynamics in diverse fields of applications such as human-gut microbiome therapies [9–13] and agroecosystem microbiome control [14–16].

**Fig. 1.**
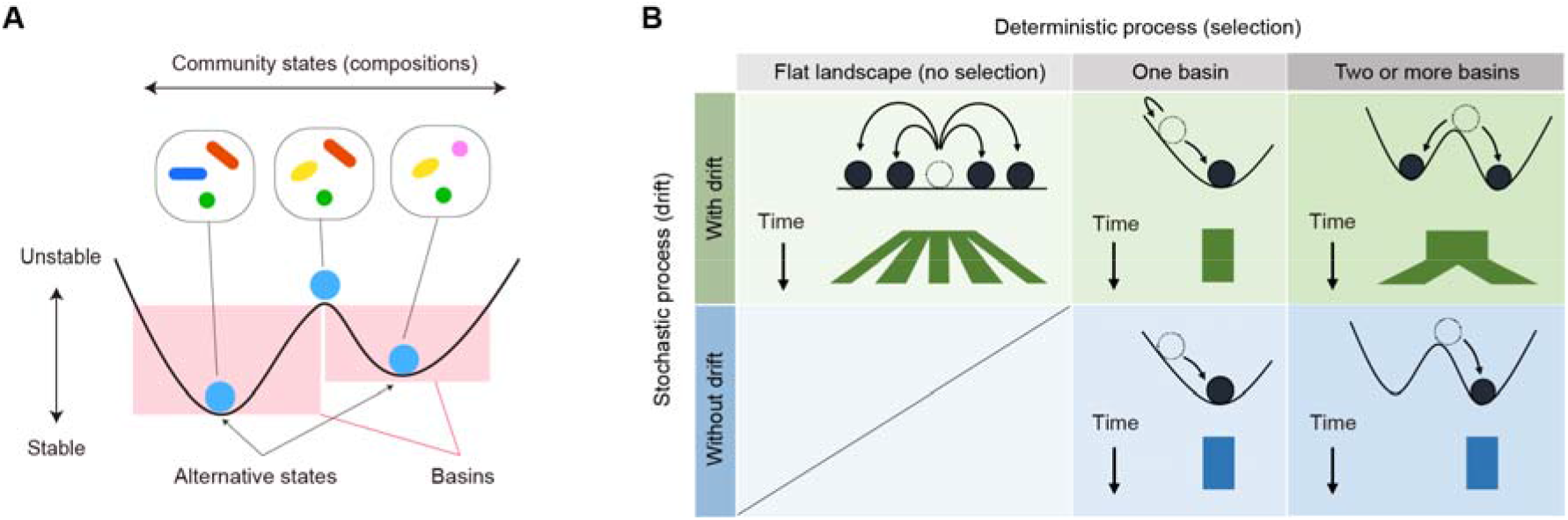
Conceptual framework of community assembly. **A** Stability landscape of community structure. Community assembly is often discussed based on the schema of “stability landscapes”, which represent the stability/instability of community states (i.e., species or taxonomic compositions). Alternative states are defined as bottom positions within basins on the stability landscapes. **B** Deterministic and stochastic processes in community assembly. Our experiment was designed to test the contributions of deterministic and stochastic ecological processes. Specifically, the results of our multi-replicate microbiome experiments are expected to depend on the presence/absence of selection and drift. Without drift (demographic fluctuations), all replicate communities will converge to a single community state (bottom panels). With drift, differentiation of community states can occur depending on the structure of stability landscapes. In particular, if there are two or more basins, differentiation into a small number of alternative community states will be observed as a consequence of both selection and drift (top right).

In empirical studies of ecological communities, consequences of deterministic and stochastic ecological processes are observable as “alternative states” of community structure [3, 17–19]. Pioneering studies on human-gut microbiomes have shown that microbiome structure can be classified into a small number of categories (i.e., “enterotypes”) despite potential numerous combinations of species or taxonomic compositions [20, 21]. The fact that only a few community states out of innumerable possible states are realized implies the presence of strong deterministic processes organizing community structure [22–24]. Meanwhile, the presence of alternative community compositions suggests that stochastic processes play some roles in the community differentiation. In other words, because no variation in community structure is expected under the assumption of strict deterministic processes, both deterministic and stochastic processes are necessary for the existence of categorizable community compositions [18, 22] (Fig. 1). Thus, empirical studies on community structure give essential insights into ecological community assembly. Nonetheless, it is basically difficult to develop detailed discussion on ecological community processes based on existing microbiome datasets because potential influence of environmental conditions on community structural patterns cannot be fully understood in observational studies.

In this respect, experimental studies with fully controlled environmental conditions are expected to provide ideal opportunities for examining ecological community processes in light of theories on alternative states [6, 8, 19]. By making a number of experimental microbial communities with defined environmental conditions, we can perform strict tests of the emergence of alternative community states. In other words, the presence of multiple reproducible states in the same experimental treatment is interpreted in the framework of selection, diversification dispersal, and drift. Despite the potential contributions of the experimental approaches to our fundamental knowledge of community assembly, few attempts (but see [25]) have been made to explore alternative states of microbiomes with tens of replications.

In this study, we examined the presence of alternative community states by performing microbiome experiments under eight nutritional (medium) conditions with 48 replicates. We constructed experimental microbiomes using a forest-soil-derived community of prokaryotes as a source microbiome and then kept the 384 microbial communities (8 treatments × 48 replicates) under a fully controlled temperature condition. The experimental system was designed to examine the roles of selection and drift in microbial community assembly. With the aid of an automated pipetting system equipped in a clean laboratory environment, we monitored changes in microbiome community compositions every two days for 12 days based on the DNA metabarcoding of 16S rRNA gene sequences. The analysis of more than 2,000 community samples then allowed us to understand how deterministic and stochastic processes could generate alternative states of ecological community structure.

## MATERIALS AND METHODS

### Terminology

In analyzing the data obtained in this study, we need to use consistent terminology to minimize the risk of confusion and misunderstanding. It is necessary to confirm the definitions of the ecological processes whose meaning can change depending on contexts. Specifically, we use the terms “selection”, “drift”, “dispersal”, and “diversification (speciation)” in ecological processes as conceptualized by Vellend [5]. Among the terms, selection represents expected changes in local community compositions resulting from differences in mean fitness between species, constituting purely deterministic processes [7]. Drift refers to demographic fluctuation that occurs regardless of among-species difference in mean fitness, forming purely stochastic processes [7]. On the other hand, dispersal itself is not a term representing stochasticity or determinism, but it merely describes processes by which organismal individuals or their propagules move between local communities. Diversification is defined as evolutionary differentiation of genetic variants and hence it represents both stochastic (e.g., nucleotide mutation) and deterministic (i.e., natural selection) phenomena [3, 8].

Another important term we need to make clear before describing our microbiome experiment is “alternative states”. In ecology, the term “alternative stable states” is often used to discuss the processes by which divergence of community compositions are caused [17, 19, 26, 27]. Meanwhile, it is generally difficult to know whether the observed community compositions are at stable states (i.e., equilibria) or they are in transient processes towards stable states [28, 29]. Stability of communities has been a central topic in the history of community ecology [30, 31]. In this study, however, we investigate divergence of community compositions irrespective of the concept of community stability. In other words, our community dataset can include information of both transient and stable states.

### Continuous-culture of microbiomes

In the microbiome experiment, the source microbiome derived from the soil of the A layer (0- 10 cm in depth) in the research forest of Center for Ecological Research, Kyoto University, Shiga, Japan (34.972 °N; 135.958 °E). After sampling, the soil was sieved with a 4-mm stainless mesh and then 5 g of the sieved soil was mixed in 100 mL PBS buffer with cycloheximide [137 mM NaCl, 8.1 mM Na_2_HPO_4_, 2.68 mM KCl, 1.47mM KH_2_PO_4_, and 200 μg/ml cycloheximide]. In this process, we added cycloheximide in order to exclude eukaryotes from the source microbiome. The source prokaryote microbiome was cultured at 22 °C for 48 hours.

We introduced the inoculum microbiome into eight types of media with different constitutions of carbon sources with 48 replicate communities per medium type (in total, 8 media × 48 replicates = 384 experimental communities; Fig. 2). To make the compositions of the media as simple as possible, we used M9 medium with minimal inorganic additives and combinations of three types of the carbon resources, specifically, glucose, leucine, and citrate as detailed in Table S1. Specifically, each of the eight media was designed based on the concentrations or presence/absence of the three carbon sources: i.e., low or high concentration of glucose (G/HG), with or without leucine (-/L), and with or without citrate (-/C). Hereafter, these medium types were designated as Medium-G (low glucose, without leucine, without citrate), GL (low glucose, with leucine, without citrate), GC (low glucose, without leucine, with citrate), GLC (low glucose, with leucine, with citrate), HG (high glucose, without leucine, without citrate), HGL (high glucose, with leucine, without citrate), HGC (high glucose, without leucine, with citrate), and HGLC (high glucose, with leucine, with citrate), respectively (Table S1).

**Fig. 2.**
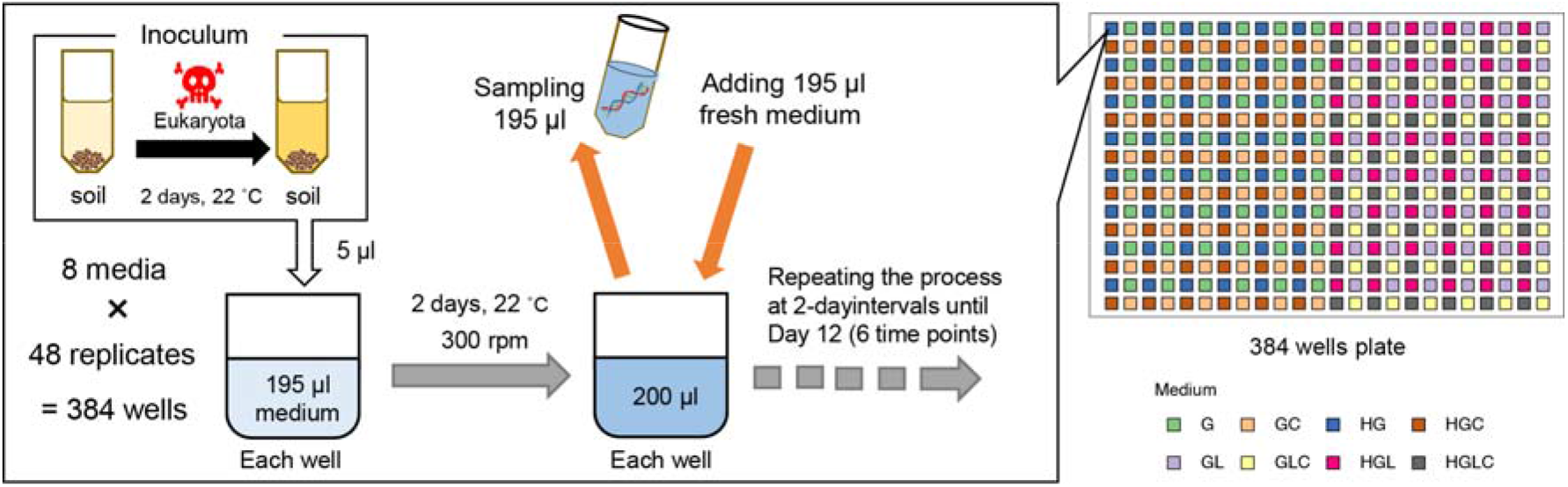
Experimental design. Laboratory culture system. A source microbiome deriving from forest soil was pre-cultured with cycloheximide under room temperature for 2 days in order to remove eukaryotes. The microbiome inoculum was then introduced into eight types of media (Table S1) with 48 replicates. A fraction of the culture fluid was sampled every 2 days and equivalent volume of fresh medium was added to the continual culture system throughout the 12-day experiment (8 media × 48 replicates × 6 time points = 2,304 community samples).

In each well of a 240 μL deep-well plate, 10 μL of the diluted source microbiome solution and 190 μL of medium were installed. Based on the quantitative amplicon sequencing detailed below, the source microbiome solution was estimated to contain 3.12 × 10^6^ DNA copies of the 16S rRNA gene (SD = 8.37 × 10^5^) (Fig. S1). Note that a previous research estimated that a single cell of *Enterobacteriaceae* bacteria has an average of 7.3 DNA copies of 16S rRNA genes (SD = 0.9; calculated with rrnDB; [32]), although variation in 16S rRNA gene copy numbers among prokaryote taxa has been known [32]. The deep-well plate was kept shaken at 200 rpm using a plate thermo-shaker BSR-MB100-4A (Bio Medical Sciences Co. Ltd., Tokyo) at 30 °C for two days. After two-days incubation, 190 μL out of the 200-μL culture medium was sampled from each of the 48 wells after mixing (pipetting) every two days for 12 days. All pipetting manipulations were performed with high precision using an automatic pipetting machine (EDR-384SR, BIOTEC Co. Ltd., Tokyo). In each sampling event, 190 μL of fresh medium was added to each well so that the total culture volume was kept constant. In total, 2,304 samples (384 communities/day × 6 time points) were collected.

Note that our study was not designed to examine historical contingency (priority effects) because the microbial species were simultaneously introduced into the experimental media. Dispersal between replicate communities were prohibited and diversification leading to speciation events was unlikely to occur in the 12-day experiment. Thus, our aim in this study was to examine how selection and drift could drive microbiome dynamics (Fig. 1).

### DNA extraction

To extract DNA from each culture sample, 5 μL of the collected aliquot was mixed with 1 μL lysozyme solution [50 mg/ml lysozyme (Sigma), 20 mM Tris-HCl (pH 8.0), 2 mM EDTA] and the mixed solution was incubated at 37 °C for 2 hours. After adding proteinase K solution [1/30 (v/v) Proteinase K (Takara), 20 mM Tris-HCl (pH 8.0), 2 mM EDTA], the aliquot was incubated at 55 °C for 3 hours and 95 °C for 10 min, The solution was then vortexed for 10 min to increase DNA yield.

We extracted DNA from the inoculum aliquot to reveal community structure of the source microbiome. Because the source inoculum was expected to include high concentrations of soil-derived compounds, which can inhibit PCR reactions, a commercial DNA extraction kit optimized for soil samples was used. Specifically, after 50 μL of the inoculum sample were incubated with 350 μL SDS buffer with proteinase K [1/30 (v/v) Proteinase K (Takara), 0.5 % SDS, 2 mM Tris-HCl (pH 8.0), 2 mM EDTA] based on the temperature profile of 55 °C for 180 min and 95 °C for 10 min. The aliquot (400 μL) was then subjected to DNA extraction with DNeasy PowerSoil Kit (Qiagen).

### PCR and DNA sequencing

For the samples of the experimental microbiomes, prokaryote 16S rRNA V4 region was PCR-amplified with the forward primer 515f fused with 3–6-mer Ns for improved Illumina sequencing quality [33] and the forward Illumina sequencing primer (5’- TCG TCG GCA GCG TCA GAT GTG TAT AAG AGA CAG- [3–6-mer Ns] – [515f] -3’) and the reverse primer 806rB fused with 3–6-mer Ns and the reverse sequencing primer (5’- GTC TCG TGG GCT CGG AGA TGT GTA TAA GAG ACA G [3–6-mer Ns] - [806rB] -3’) (0.2 μM each). The buffer and polymerase system of KOD One (Toyobo) was used with the temperature profile of 35 cycles at 98 °C for 10 s, 55 °C for 5 s, 68 °C for 1 s. To prevent generation of chimeric sequences, the ramp rate through the thermal cycles was set to 1 °C/sec [34]. Illumina sequencing adaptors were then added to respective samples in the supplemental PCR using the forward fusion primers consisting of the P5 Illumina adaptor, 8-mer indexes for sample identification [35] and a partial sequence of the sequencing primer (5’- AAT GAT ACG GCG ACC ACC GAG ATC TAC AC - [8-mer index] - TCG TCG GCA GCG TC -3’) and the reverse fusion primers consisting of the P7 adaptor, 8-mer indexes, and a partial sequence of the sequencing primer (5’- CAA GCA GAA GAC GGC ATA CGA GAT - [8-mer index] - GTC TCG TGG GCT CGG -3’). KOD One was used with a temperature profile of 8 cycles at 98 °C for 10 s, 55 °C for 5 s, 68 °C for 5 s (ramp rate = 1 °C/s). The PCR amplicons of the samples were then pooled after a purification/equalization process with the AMPureXP Kit (Beckman Coulter). Primer dimers, which were shorter than 200 bp, were removed from the pooled library by supplemental purification with AMPureXP: the ratio of AMPureXP reagent to the pooled library was set to 1 (v/v) in this process. This library was further purified with E-gel SizeSelect 2 (Invitrogen) and then ca. 440-bp DNA fragments was selectively obtained. The sequencing libraries were processed in an Illumina Miseq sequencer [271 forward (R1) and 31 reverse (R4) cycles; 20 % PhiX spike-in].

For the source microbiome sample, the prokaryote 16S rRNA V4 region was amplified as well. To estimate concentrations of 16S rRNA genes included in the inoculum, a quantitative amplicon sequencing platform was applied by introducing five “standard DNA” fragments with controlled concentrations to the PCR master mix solution of the first PCR process as detailed elsewhere [36, 37]. The standard DNAs were used for the *in-silico* calibration of 16S rRNA gene concentrations in the target sample after sequencing as detailed in the previous study [36–38].

### Bioinformatics

In total, 25,284,304 sequencing reads were obtained with the Illumina sequencing. The raw sequencing data obtained in the Illumina sequencing were converted into FASTQ files using the program bcl2fastq 1.8.4 distributed by Illumina. The output FASTQ files were then demultiplexed with the program Claident v0.2. 2018.05.29. The sequencing reads were subsequently processed with the program DADA2 [39] v.1.18.0 of R 3.6.3 to remove low-quality data. The molecular identification of the obtained amplicon sequence variants (ASVs) was performed based on the naive Bayesian classifier method [40] with the SILVA v.132 database [41]. Based on the *in-silico* calibration the quantitative amplicon sequencing data [36, 37], the number of DNA copies in 10 μL of source microbiome (inoculum) was estimated to be 3.12 × 10^6^ copies as mentioned above (Fig. S1).

### Community-level diversity

The rarefaction curves representing relationship between the number of sequencing reads and the number of ASVs were drawn using the vegan 2.6.4 package [42] of R. In the sequencing of experimental culture samples, the diversity of microbial ASVs reached plateaus along the axis of the number of sequencing reads (Fig. S2). Given the rarefaction curves, the dataset was rarefied to 5,000 reads per sample with the rrarefy function of the R vegan package. Of the 2,304 samples (8 treatments × 48 replicates × 6 time points), 2,250 samples with more than 5,000 reads were used in the following pipeline. After screening for the replicate communities for which sequencing data were available for all the six time points, 2,094 samples were subjected to the following statistical analyses. In total, 718 prokaryote ASVs belonging to two kingdoms, 19 phyla, 32 classes, 74 orders, 89 families, and 115 genera were detected.

For each sample, two types of *α*-diversity indices, ASV richness and Shannon-Wiener diversity, were calculated. We then compared *α*-diversity among each medium using Student’s *t*-test corrected by false discovery rate (FDR) by Benjamini-Hochberg method (Table S2).

### Overview of the community structure

To visualize the diversity of the prokaryote community structure, an analysis of non-metric multidimensional scaling (NMDS) was performed based on the Bray-Curtis-metric of *β*- diversity. Likewise, to examine the dependence of community structure on medium conditions, a series of permutational multivariate analysis of variance (PERMANOVA) [43] was performed with 50,000 permutations. In each PERMANOVA model, glucose concentration (high or low; df = 1), the presence/absence of leucine (df = 1), or the presence/absence of citrate (df = 1) was included as the explanatory variable. An additional model including all the medium conditions and interactions between them (df = 7) was examined as well. Each PERMANOVA model was supplemented by permutational analysis of dispersion (PERMDISP; [44]) for potential effects of the medium condition on dispersion in community structure. The NMDS, PERMANOVA, and PERMDISP were performed for each dataset of the ASV-, genus-, and family-level compositions of prokaryote communities using the vegan 2.6.4 package [42] of R.

### Community structural differentiation

To examine the differentiation of community compositions among replicate samples, the Bray-Curtis metric of *β*-diversity was calculated for respective pairs of samples collected on the same day in each of the same treatments (medium conditions). Note that the Bray-Curtis metric of *β*-diversity could range from 0 (identical community structure) to 1 (no overlap of microbes). A histogram of the community structural difference was shown for each day in each experimental treatment. If a small number of alternative states of community structure exist under a medium condition, there can be some peaks of community structural difference within a histogram. The analysis was performed for each dataset of the ASV-, genus-, and family-level compositions of the prokaryote communities.

### Potential differentiation in community-level functions

To examine potential differentiation of ecosystem functions among the replicate communities, we inferred the community-level functional profiles based on the phylogenetic estimation of gene repertoires with PICRUSt2 [45]. The gene repertoire data were then subjected to NMDS, PERMANOVA, and PERMDISP based on Bray-Curtis *β*-diversity. In total, the relative functional compositions of 392 pathways were examined as input data. As examined in the previous section about community structural differentiation, histograms of Bray-Curtis *β*- diversity were drawn at respective time point within respective treatments based on the inferred gene repertoires.

## RESULTS

### Overview of the community structure

In each of the experimental treatments (medium conditions), taxonomic richness and Shannon’s diversity index at the ASV, genus, and family levels decreased through time (Fig. S3-4). The α-diversity indices varied depending also on medium conditions (Table S2). The addition of leucine or citrate, for example, significantly increased α-diversity of the experimental microbiomes (FDR: q < 0.01) (Fig. 3A; see Table S2).

**Fig. 3.**
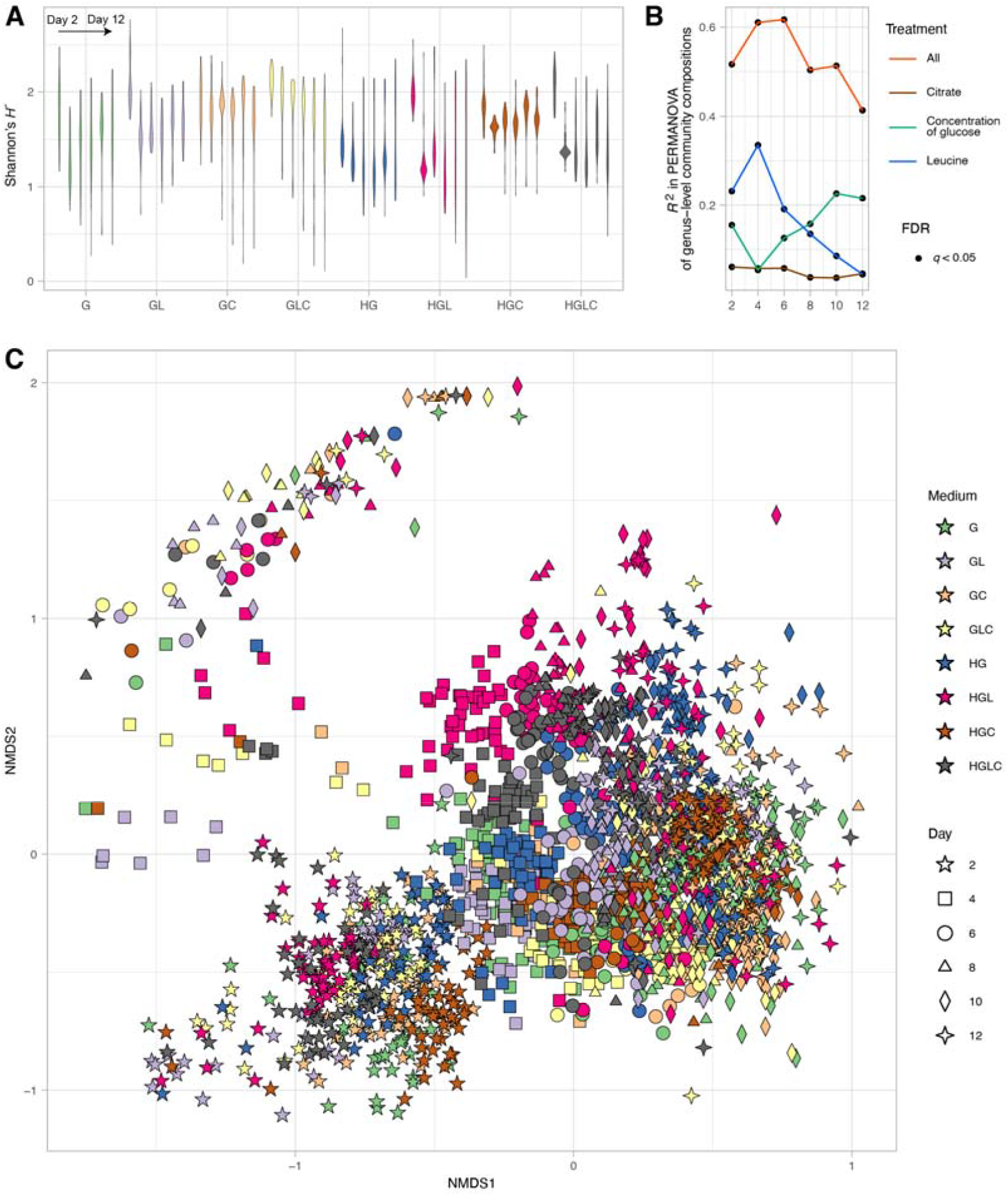
Overview of the community structure. **A** *α*-diversity of the communities. In each experimental treatment (medium condition), Shannon’s diversity index for ASV-level community compositions is shown for each day. The results of Student’s *t*-test are shown in Table S2. **B** Dependence of community structure on medium conditions. In each PERMANOVA model of ASV-level community compositions, glucose concentration (high or low; df = 1), the presence/absence of leucine (df = 1), or the presence/absence of citrate (df = 1) was included as the explanatory variable. An additional model including all the medium conditions and interactions between them (df = 7) was examined as well. The coefficient of determination (*R^2^*) is shown for each day. **C** Overview of the community structure. The ASV- level community compositions are shown along the axes of nonmetric multidimensional scaling (NMDS) (stress = 0.157).

At the genus level, the microbiomes were dominated by the four genera, *Klebsiella*, *Raoultella*, *Pseudomonas,* and *Cedecea*, although there were substantial variation in the balance of these taxa among the medium conditions examined (Fig. 4). For example, in the media containing citrate (Medium-GC, GLC, HGC and HGLC), the relative abundance of *Citrobacter* was higher than in other medium conditions. In the media containing leucine (Medium-GL, GLC, HGL and HGLC), *Cedecea* were more abundant than in other medium conditions. Moreover, within each of the medium conditions, *Serratia* became dominant only in some replicate communities (Fig. 4).

**Fig. 4.**
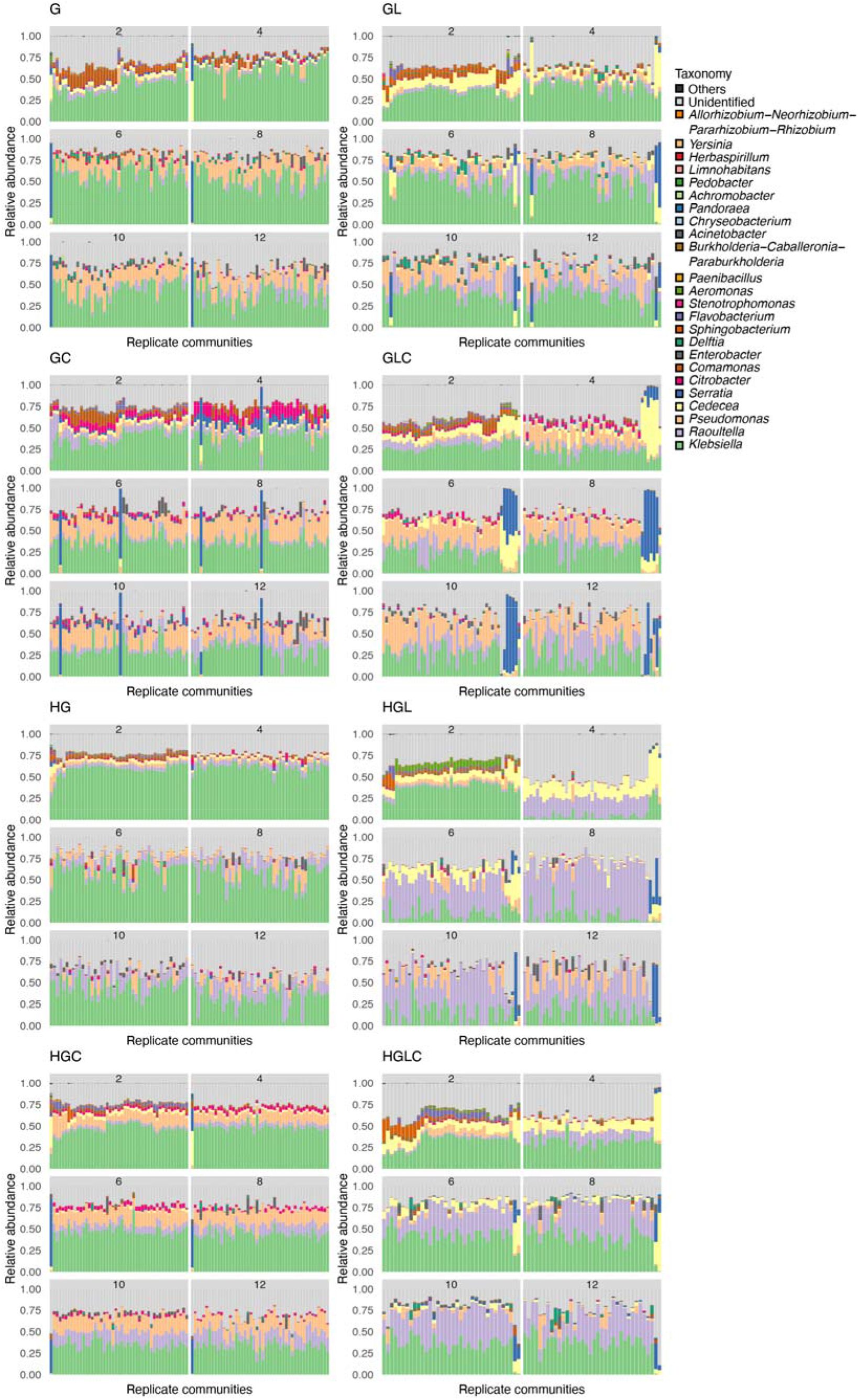
Variation in community structure among replicate samples. For each replicate community in each experimental treatment, changes in genus-level community compositions (relative abundance) are shown. The numbers shown at the top of the bar plots refer to time points (days). The replicate samples were ordered based on unweighted pair group method with arithmetic mean (UPGMA) analyses performed on Day 2 for respective experimental treatments. The order of replicate communities on successive days is the same as that on Day 2.

At the family level, the communities were characterized by *Enterobacteriaceae*, *Pseudomonadaceae*, and *Comamodaceae*, with *Enterobacteriaceae* being particularly dominant (Fig. S6). *Comamodaceae* tended to appear at high proportions in the media with high concentration of glucose. The substantial among-replicate variation in community compositions observed at the ASV- and genus-level analyses was evident as well in the family-level analysis. Specifically, in the media containing leucine (medium-GL, GLC, HGL and HGLC), *Yersiniaceae*, which included *Serratia*, was dominated only in some replicate communities within each experimental treatment.

At all time points, differences in community compositions were significantly explained by differences in medium (Figs. 3B, S7-8 and Table S3). The results also indicated that the overall variance explained by medium conditions decreased from Day 4 to 12 (Figs. 3B and S7). Among the components of the medium conditions, concentrations of glucose had the largest contributions to community structure at the family level (Fig. S7). Meanwhile, the presence/absence of leucine had the highest impacts on the ASV- and genus-level community structure (Fig. S7). The community structure within the NMDS plot distributed depending on both time points and medium conditions (Fig. 3C). The PERMDISP indicated that the concentration of glucose and the presence/absence of citrate greatly influenced dispersion of family-level community structure among samples (Fig. S8 and Table S4). In contrast, the presence/absence of leucine was the major determinant of community structural dispersion at the genus- and ASV-level (Fig. S8).

### Community structural differentiation

In some medium conditions (experimental treatments), large community structural difference among replicate samples were observed. For example, in the Medium-GLC treatment, substantial difference in community compositions were observed among replicate samples at the ASV, genus, and family levels, and these differences seemingly increased until Day 10 (Figs. 5, S9 and S10). The community compositions seemed to be classified into some categories within the NMDS plot (Figs. 5, S9 and S10) due to the dominance of *Serratia* in some, but not all, replicate samples (Fig. 4). In fact, the histogram of community structural difference between samples (i.e., Bray-Curtis *β*-diversity between pairs of replicate samples) showed bimodal or multi-modal distributions from Day 2 to Day 10, although eventually displayed a uniform distribution on Day 12 (Figs. 6, S11 and S12). Such bimodal or multimodal distributions of *β*-diversity were observed in all the eight medium conditions (Fig. 6), although among-replicate differentiation in community structure on the NMDS surface was conspicuous in some treatments (Medium-G, GL, GLC, HGL, HGC, and HGLC) but not in others (Medium-GC and HG) (Fig. 5).

**Fig. 5.**
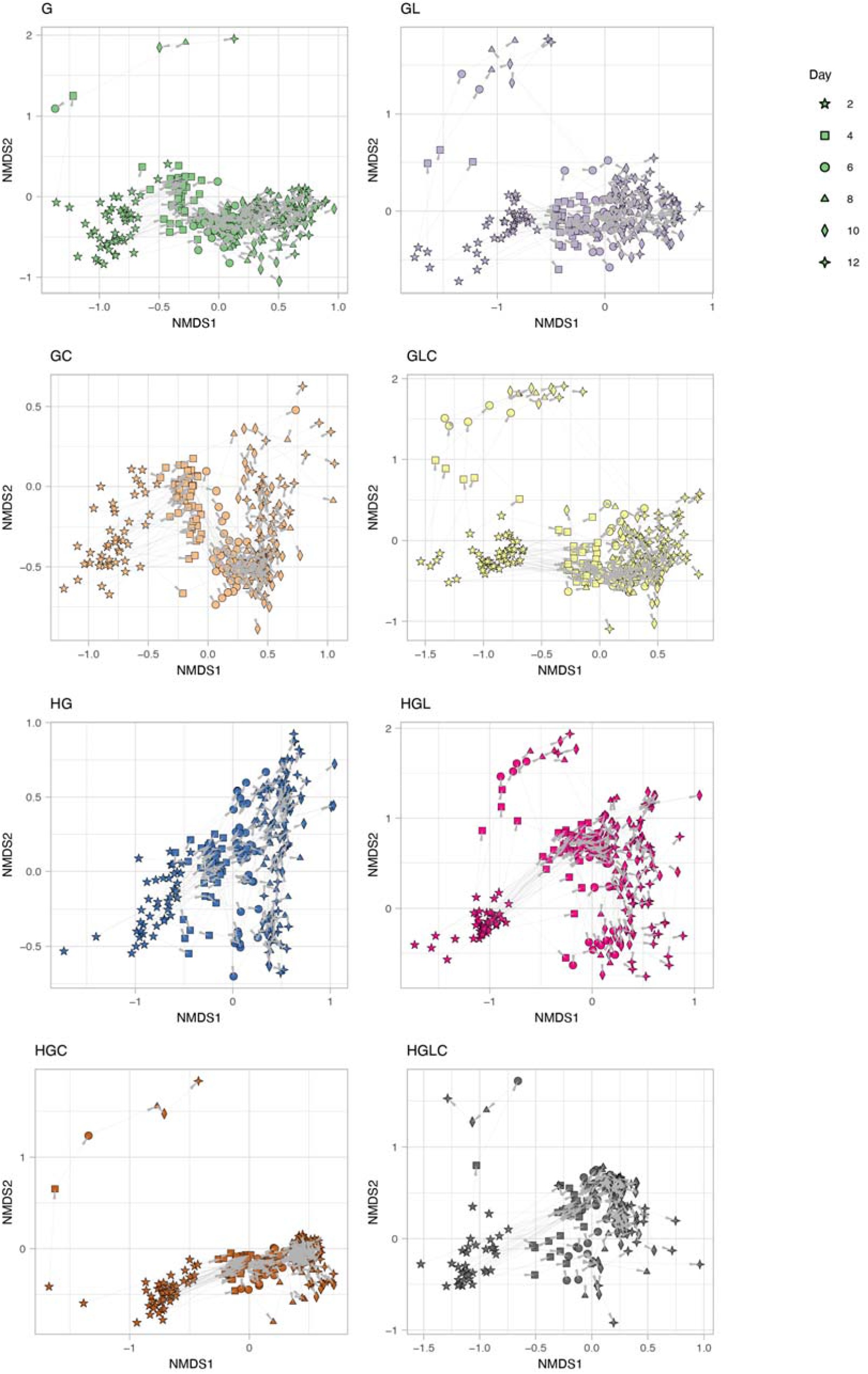
Time-series changes in community structure. For each replicate community in each experimental treatment, time-series changes in ASV-level community structure are shown with arrows on the NMDS surface defined in Figure 3C.

**Fig. 6.**
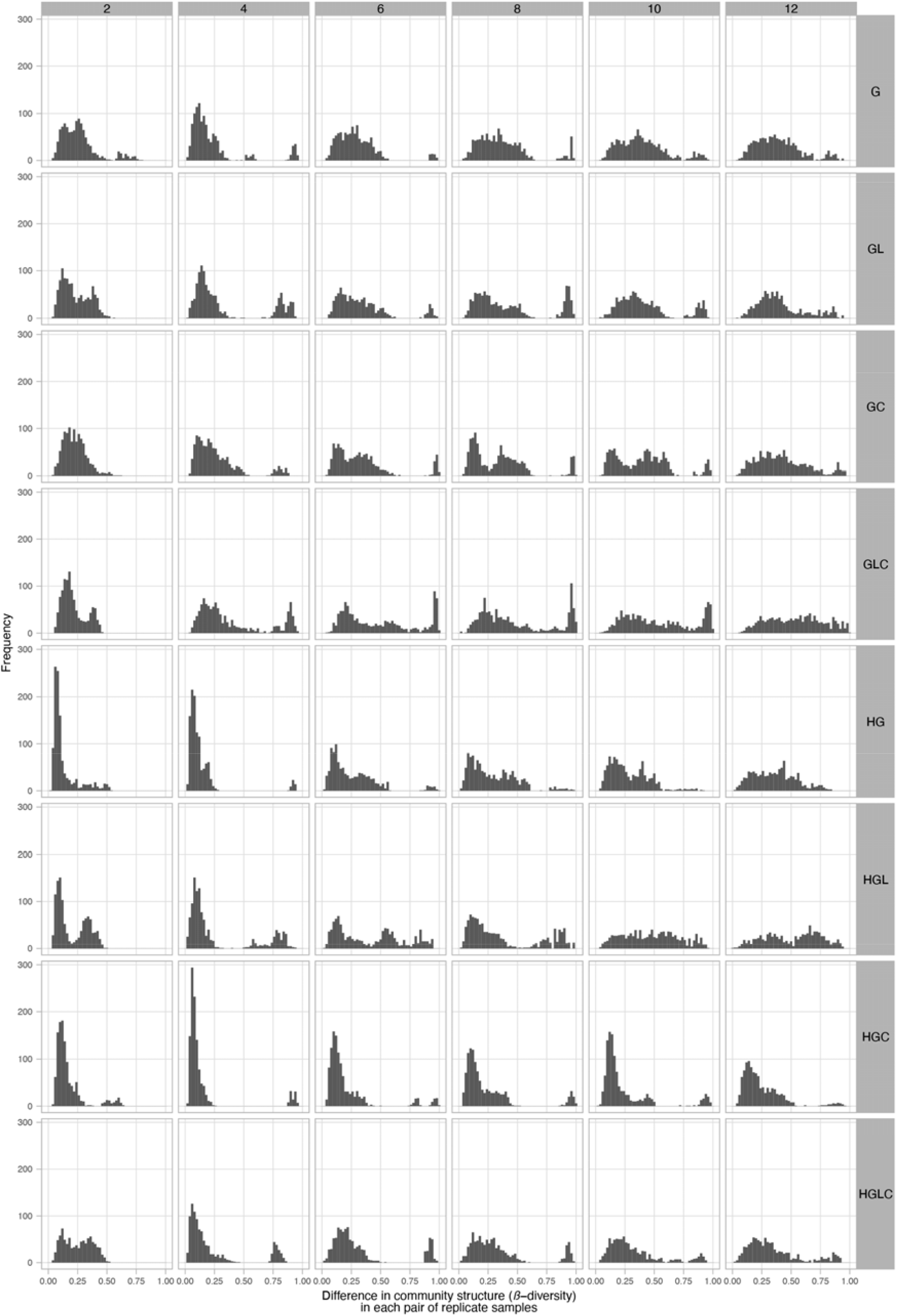
Histograms of community structural differentiation. For each experimental treatment, difference in ASV-level community structure (Bray-Curtis *β*-diversity) between replicate communities is shown as a histogram for each day. The numbers shown at the top of the histograms refer to the time points (days). The bi-modal or multi-modal distributions within these histograms suggest the presence of alternative community states.

Within the deep-well plate used in the experiment, substantially different community compositions were observed between adjacent replicate wells in each experimental treatment (Fig. S15, 16). This fact suggests that potential fine-scale heterogeneity of temperature or humidity within the small culture plate does not fully explain the observed among-replicate divergence of community structure.

### Potential differentiation in community-level functions

As observed in the above analyses on the community compositions, functional profiles of the community samples significantly differed depending on the medium conditions (Fig. 7A-B). Moreover, among-replicate differentiation was evident in the functional profile dataset (Fig. 7C-D), although the extent of such differentiation varied among medium conditions (Fig. S13-14).

**Fig. 7.**
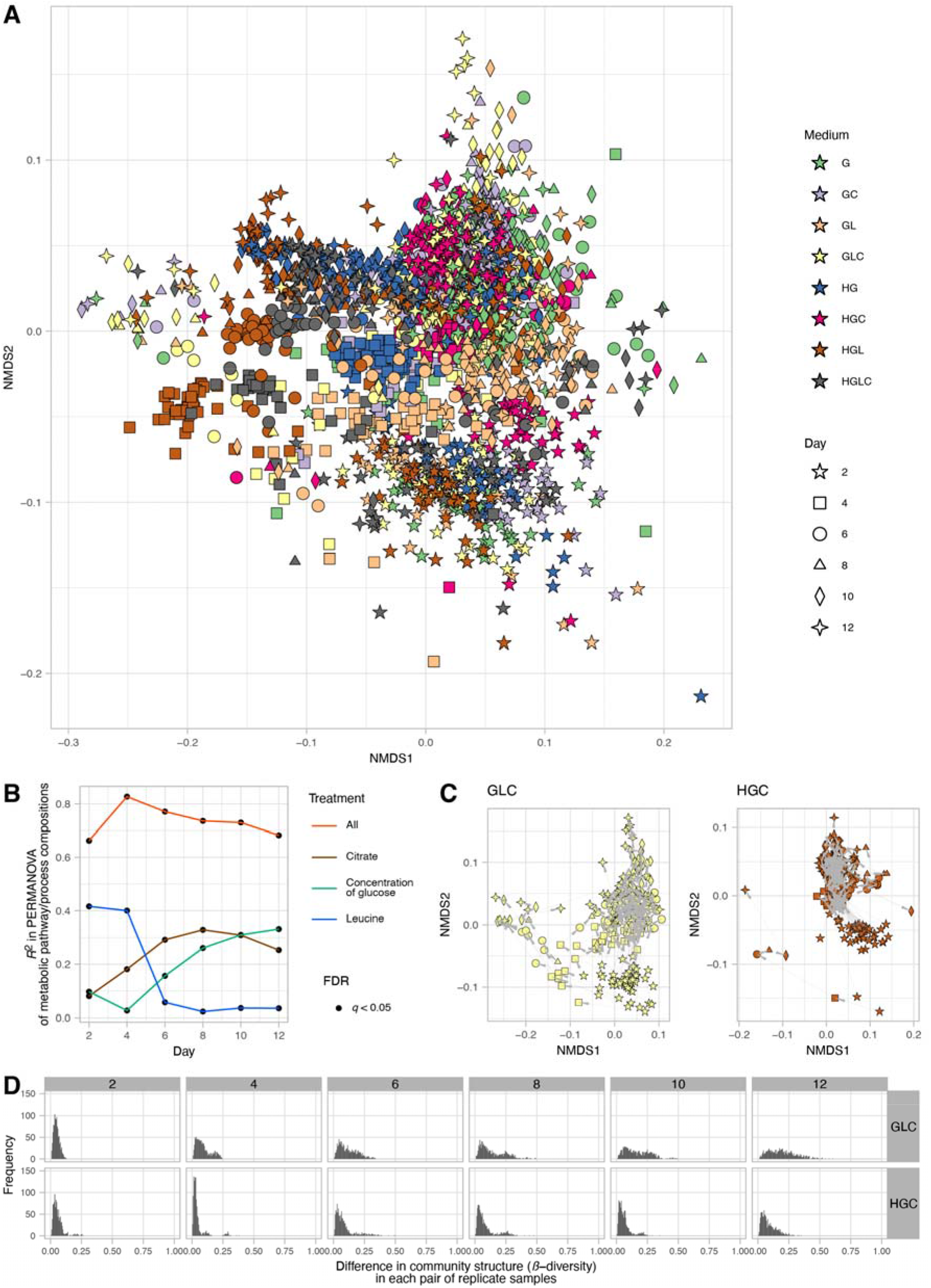
Functional profiles of the microbiomes. **A** Overview of functional compositions. Based on a phylogenetic inference of metabolic pathway compositions (a reference-genome analysis of constituent bacteria based on PICRUSTs2), community samples are plotted on a NMDS surface (stress = 0.125). **B** Dependence of community functional profiles on medium conditions. In each PERMANOVA model of metabolic pathway compositions, glucose concentration (high or low; df = 1), the presence/absence of leucine (df = 1), or the presence/absence of citrate (df = 1) was included as the explanatory variable. An additional model including all the medium conditions and interactions between them (df = 7) was examined as well. The coefficient of determination (*R^2^*) is shown for each day. **C** Time-series changes in community functional profiles. The results on two experimental treatments (Medium GLC and HGC) are shown. See Fig. S13 for full results. **D** Histograms of community functional differentiation. The results on two experimental treatments (Medium GLC and HGC) are shown. See Fig. S14 for full results.

## Discussion

Based on the experimental design with many replicates in each of the multiple treatments, we examined how microbial community assembly was driven by deterministic and stochastic processes. Hereafter, we discuss respective components of those ecological processes in the framework outlined in Fig. 1.

In terms of deterministic ecological processes, the importance of selection has been discussed for decades in microbiology [7, 8, 46–50]. Indeed, in our study, the community structure differed remarkably depending on the medium conditions (Fig. 3), indicating that selection [8, 49] operated in the community processes of the experimental microbiomes. For example, *Citrobacter*, which has the ability to metabolize citrate, occurred almost exclusively in the media containing citrate (Fig. 4). Thus, selection can be regarded as a fundamental mechanism determining community structure.

In addition to selection, drift was shown to organize the community structure of the experimental microbiomes. We found that community compositions could diverge into a small number of reproducible alternative states even under the same environmental (medium) conditions (Fig. 3-6). This finding indicates that selection and drift could collectively generate alternative states of community structure in microbiome dynamics (Fig. 1). On a stability landscape of community structure, divergence into a small number of reproducible states never occur without selection (Fig. 1). In addition, stochastic processes like drift (demographic fluctuations) are necessary for such divergence because without stochasticity, identical consequences are always expected. Meanwhile, careful interpretation is required when we discuss the stochastic processes caused by drift. In our experiment, drift might influence the community assembly not exclusively at the (pre-)colonization stage (i.e., stochastic sampling effects in the pipetting of inoculum microbiomes) but also at the post-colonization stage (i.e., “random walk” fluctuation of community compositions). The relative contributions of pre-colonization and post-colonization stochastic events need to be examined in future studies. In our present experiment, 3.12 × 10^6^ DNA copies of the 16S rRNA gene were introduced into each replicate community at the pre-colonization stage, while 10 μL out of 200 μL of culture media was left for successive time points (i.e., 5 % bottleneck in each transition between time points) at the post-colonization stage. Experiments comparing multiple source microbiome density and multiple post-colonization bottleneck levels will provide further crucial knowledge of how drift can drive community dynamics leading to alternative states.

In our results, community structural differences evident at the ASV or genus levels (Fig. 3-6) were less conspicuous at the family level (Fig. S6, S9 and S11) partly due to the dominance of bacteria belonging to Enterobacteriaceae. Such convergence of community structure at higher taxonomic levels may stem from redundancy in functional compositions of microbial communities as has been reported in previous studies [9, 51, 52]. However, in some experimental treatments (e.g., Medium-GLC), substantial divergence of taxonomic compositions among replicate communities was evident even at the family level (Fig. S6). Thus, substantial functional differentiation could occur through the emergence of alternative states of microbiome structure.

In fact, a reference-genome-based analysis suggested that divergence into alternative community structure could entail functional differentiation of microbial communities. The inferred community-level gene repertoires were differentiated into some clusters within each experimental treatment (Fig. 7C), resulting in bimodal or multi-modal distributions of pairwise community dissimilarity (Fig. 7D). This observation is of particular interest because knowledge of the processes driving divergence into functionally different microbiome states is essential in diverse fields of applied sciences. In medicine, for example, the structure of human gut microbiomes has been classified into some categories (i.e., enterotypes) differing in associations with host human health [20, 21]. Furthermore, recent studies have suggested that fish-associated and plant-associated microbiomes can be classified into several community compositional types potentially differing in physiological impacts on host organisms [14, 53]. To extend the discussion on functional divergence of microbiomes, the genomic information of the microbes constituting microbiomes need to be enriched with shotgun metagenomic analyses [54–56].

From a detailed inspection of the experimental results, we found that shifts between alternative states could occur in microbial community dynamics. While difference in community structure were expanded from Day 2 to Day 10, transitions between alternative states seemed to occur within the NMDS plots in some experimental treatments (Medium-GL, GLC, and HGL; Fig. 5). This observation illuminates the ecological theory that transitions between alternative stable states are possible if demographic fluctuations of microbial populations within the communities are large enough to cross the “boundaries” splitting basins of stability landscapes [26, 27]. Alternatively, the observed dynamics may be interpreted as transient dynamics towards a large basin within a stability landscape [19, 29]. Although it is notoriously difficult to distinguish alternative transient states from alternative stable states based on current frameworks of empirical datasets, further feedback between theoretical and empirical investigations will promote our understanding of large shifts in community structure [38].

The fact that patterns in the divergence into alternative states differed among environmental conditions give significant implications for the “controllability” of microbiomes. We found that the number and structure of alternative states differed depending on medium conditions (Fig. 5). This result suggests that the shapes of underlying stability landscapes, which are formed by selection, can be changed by the addition of specific chemicals to the microbial ecosystems. Such changes in stability landscape structure have been intensively discussed in theoretical ecology [27] but explored in a few empirical studies [57, 58]. Thus, this study indicates that microbiome experiments under a series of environmental conditions provide ideal opportunities for investigating how microbiome structure can be managed by changing the structure of background stability landscapes based on the manipulation of environmental conditions.

While the experiment using field-collected source microbiomes allowed us to explore broad state space of possible community structure, experiments based on explicitly defined sets of microbial species will provide complementary insights. A more mechanism-based understanding may become possible by conducting similar experiments on multispecies systems with known genomes and metabolic pathways. In this respect, experiments on synthetic communities (SynCom) [59–61] are expected to promote reductionistic understanding of microbial community processes [62, 63]. With the aid of genome-based metabolic modeling [64–66], for example, potential consequences of competitive and facilitative interactions between microbial species may be inferred at the community level [56]. For further understanding of microbiome assembly, it is also important to apply empirical frameworks for describing complex time-series processes of ecological communities [38, 67, 68] in light of shifts between alternative states [38]. Interdisciplinary studies based on microbiology, genomics, and theoretical ecology will reorganize our knowledge of the stability and dynamics of microbiomes.

## DATA AVAILABILITY

The 16S rRNA gene sequence data are available from the DNA Data Bank of Japan (DDBJ; accession number, Bioproject PRJDB15353) [to be released after acceptance of the paper]. The microbial community data and all the R scripts used for the statistical analyses are available at the GitHub repository (https://github.com/Ibuki-Hayashi/deterministic-and-stochastic-processes-generating-alternative-states) [to be released after acceptance of the paper].

## ACKNOWLEDGEMENTS

This work was financially supported by NEDO Moonshot Research and Development Program (JPNP18016) and JST FOREST (JPMJFR2048) to HT.

## AUTHOR CONTRIBUTIONS

IH and HT designed the work. IH performed the experiments. IH and HF analyzed the data. IH and HT wrote the paper with HF.

## COMPETING INTERESTS

The authors declare no competing interests.

## Notes

### Competing Interest Statement

The authors have declared no competing interest.

