## Supplementary Information for "Deterministic and stochastic processes generating alternative states of microbiomes"

**This PDF file includes:**

Supplementary Figures S1-16

**Supplementary Information included in a separate file:**

Supplementary Tables S1-4

### SI Figures

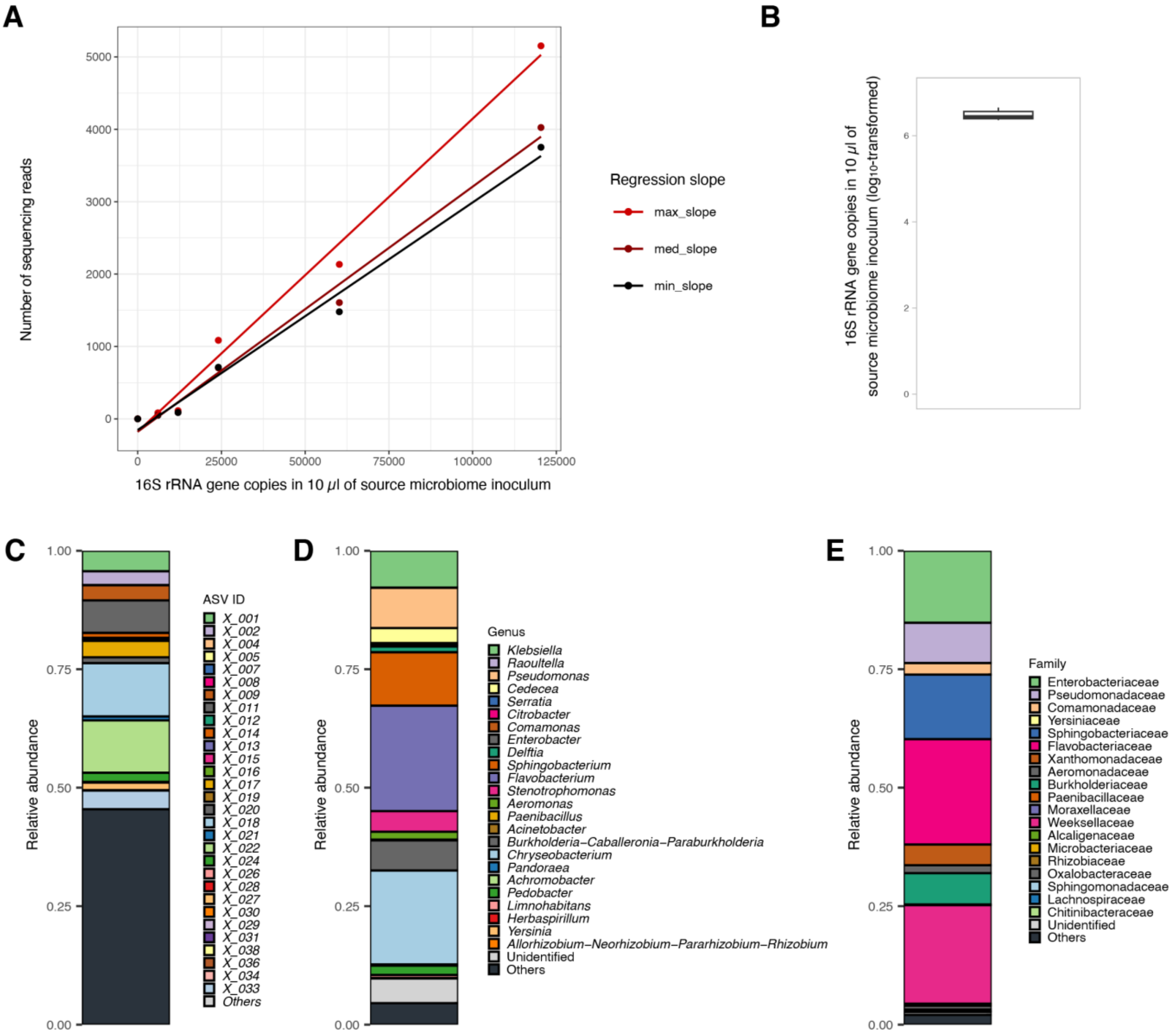


**Fig. S1 | Overview of the source microbiome. A** Calibration of 16S rRNA gene copy concentrations in the source microbiome. To estimate concentrations of 16S rRNA genes included in the source microbiome inoculum, a quantitative amplicon sequencing platform was applied by introducing five "standard DNA" fragments with controlled concentrations to the PCR master mix solution as detailed elsewhere [1, 2]. The standard DNA fragments differed in their concentrations, subjected to the *in-silico* calibration of 16S rRNA gene concentrations in the target sample after sequencing [1–3]. Correlation between 16S rRNA gene copy concentrations and the number of standard-DNA sequencing reads is shown. Among the seven replicates of source microbiome samples subjected to the calibration (left panel), the sample with the highest and lowest correlation coefficients are shown in red and black, respectively (right panel). **B** Estimated 16S rRNA gene copy concentrations. The estimated concentrations of the prokaryote 16S rRNA gene are shown as a boxplot. **C** ASV-level composition of the source microbiome. **D** Genus-level composition of the source microbiome. **E** Family-level composition of the source microbiome.

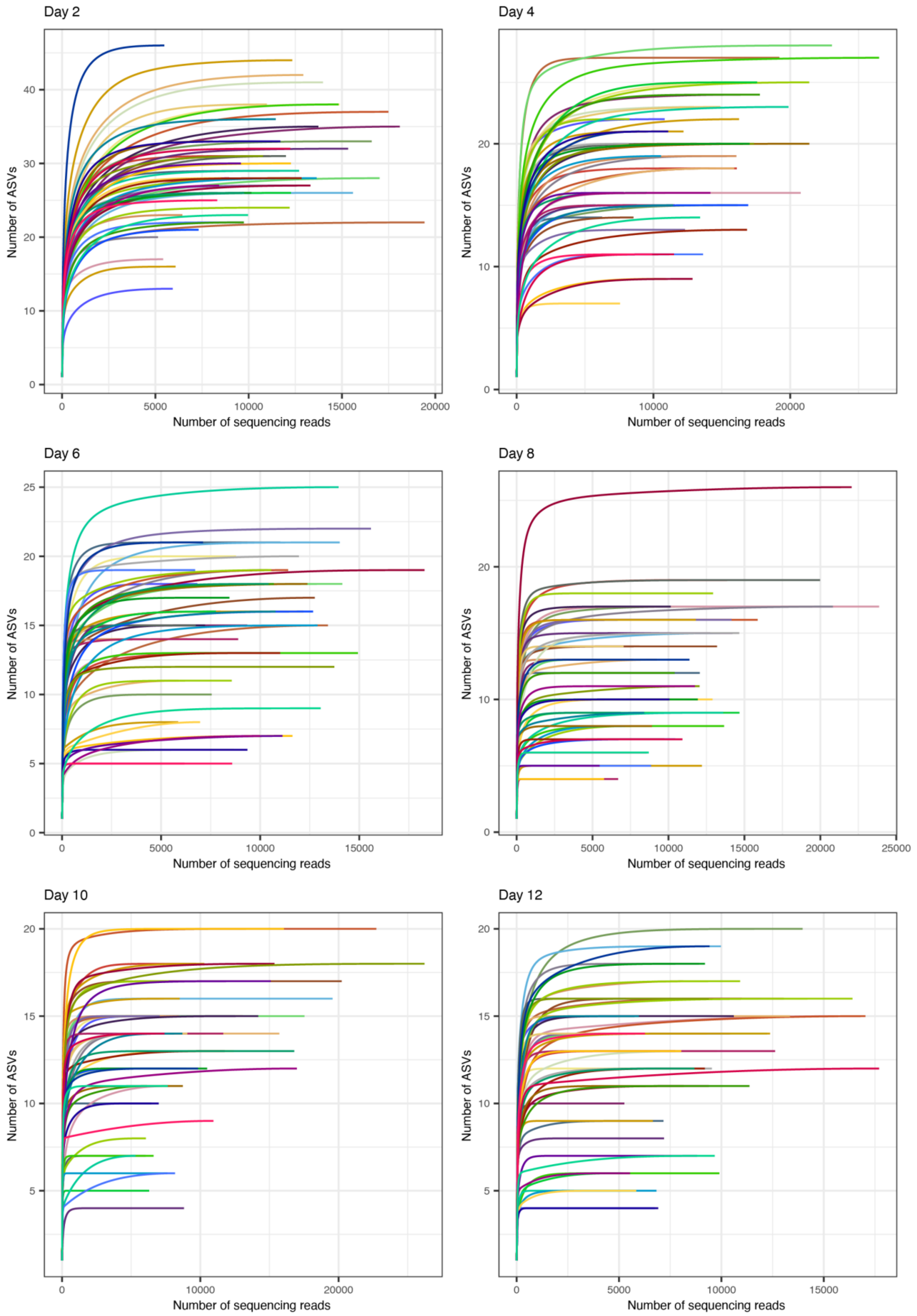
Day 2
Day 4
Day 6
Day 8
Day 10
Day 12
Number of ASVs
Number of sequencing reads

**Fig. S2 | Rarefaction curves of ASV richness.** Relationship between the number of sequencing reads and the number of detected prokaryote ASVs is shown for each day. In each panel (day), 50 samples randomly selected from the pool of the samples with 5,000 or more sequencing reads.

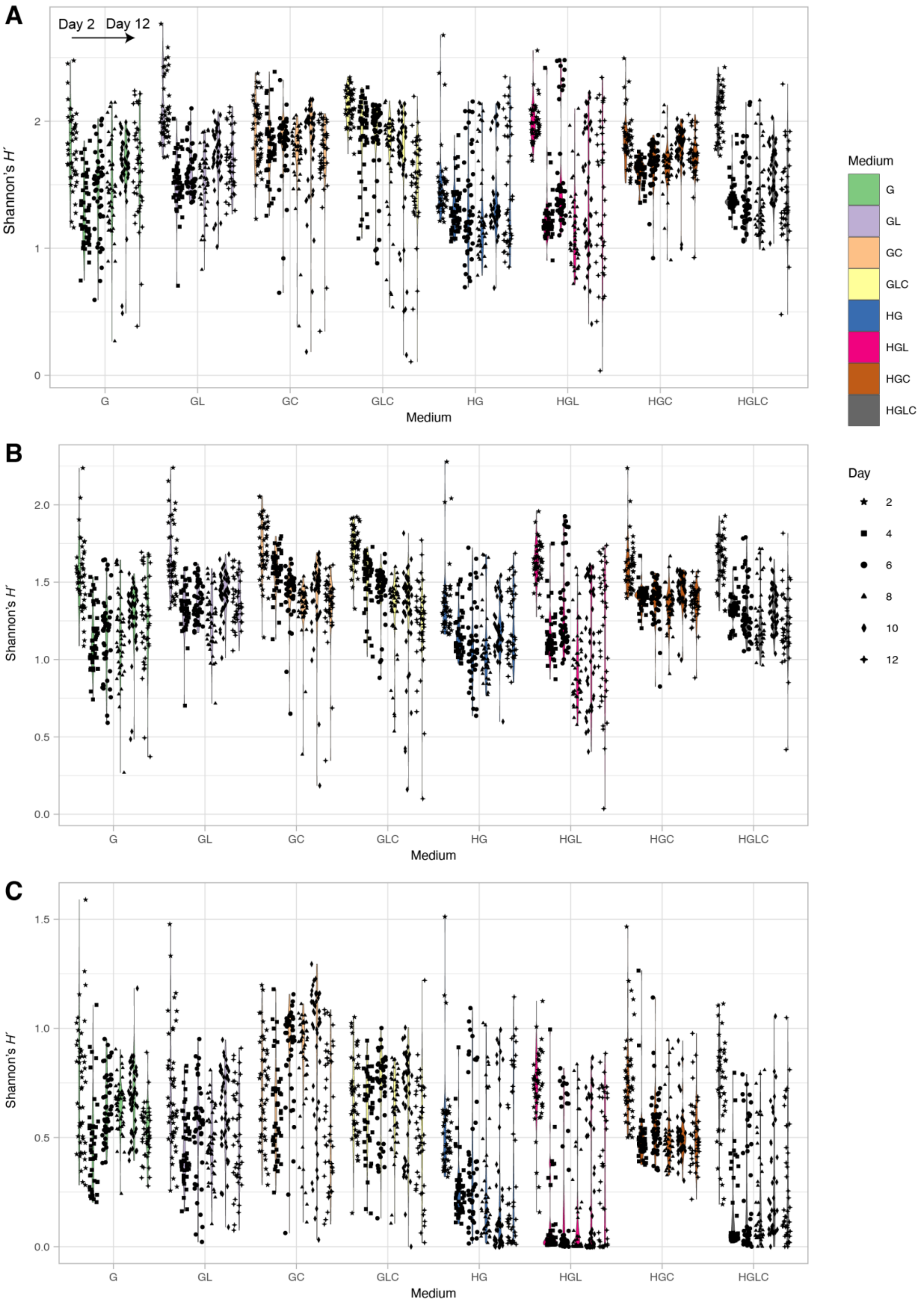


**Fig. S3 | Shannon's diversity of the community samples. A** $\alpha$-diversity of the communities. For each experimental treatment (medium condition), Shannon's diversity index for ASV-level community compositions is shown for each day. The results of Student's $t$-test are shown in Table S2. **B** $\alpha$-diversity at the genus level. **C** $\alpha$-diversity at the family level.

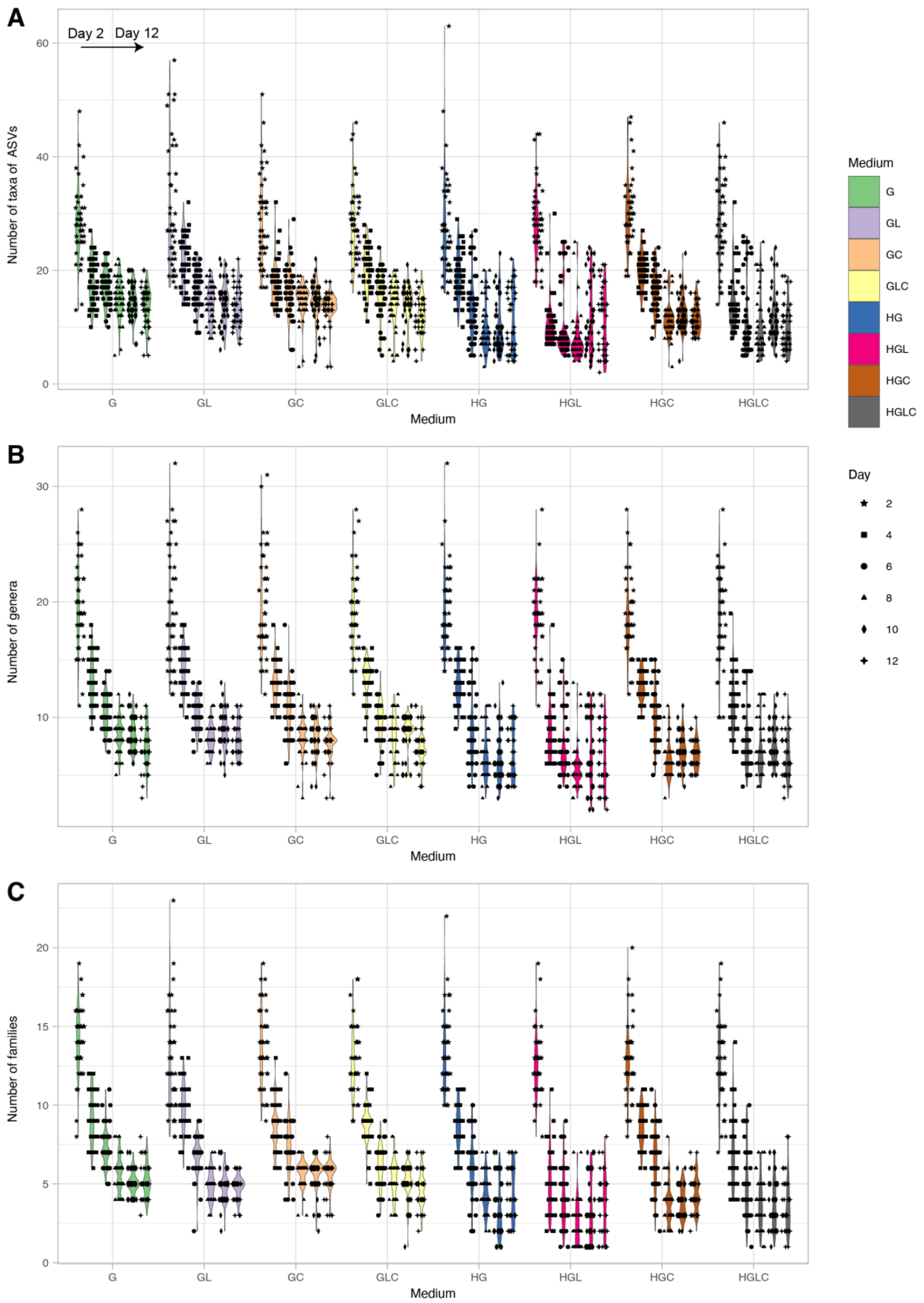


**Fig. S4 | ASV/taxonomic richness of the community samples. A** ASV-level richness. For each experimental treatment (medium condition), the number of detected prokaryote ASVs is shown for each day. The results of Student's *t*-test are shown in Table S2. **B** Taxonomic richness at the genus level. **C** Taxonomic richness at the family level.

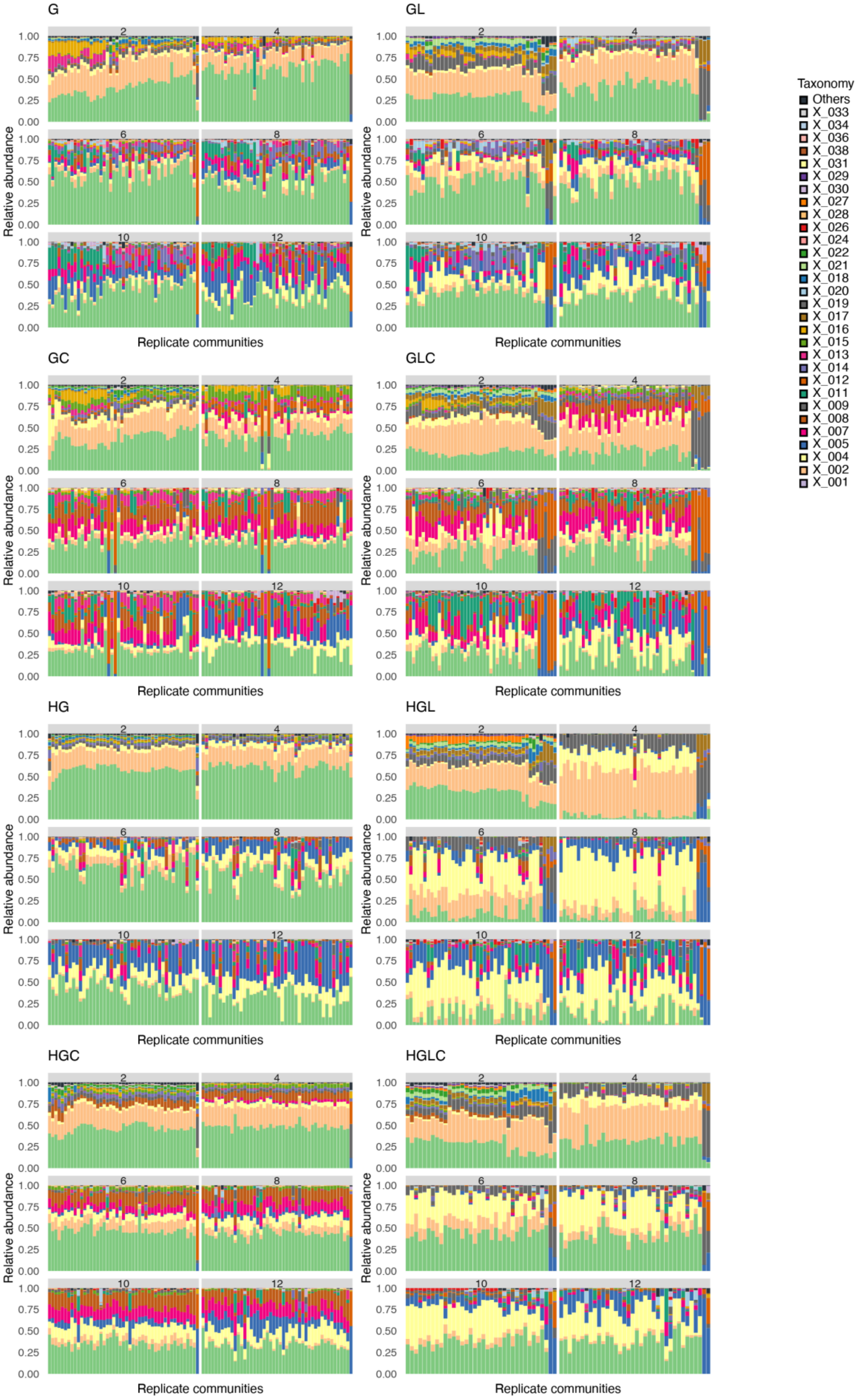
G
GL
GC
GLC
HG
HGL
HGC
HGLC
Relative abundance
Replicate communities
Taxonomy
Others
X_033
X_034
X_036
X_038
X_031
X_029
X_030
X_027
X_028
X_026
X_024
X_022
X_021
X_018
X_020
X_019
X_017
X_016
X_015
X_013
X_014
X_012
X_011
X_009
X_008
X_007
X_005
X_004
X_002
X_001

**Fig. S5 | Variation in community structure among replicate samples (ASV level).** For each replicate community in each experimental treatment, changes in ASV-level community compositions (relative abundance) are shown. The numbers shown at the top of the bar plots refer to time points (days). The replicate samples were ordered based on unweighted pair group method with arithmetic mean (UPGMA) analyses performed on Day 2 for respective experimental treatments. The order of replicate communities on successive days is the same as that on Day 2.

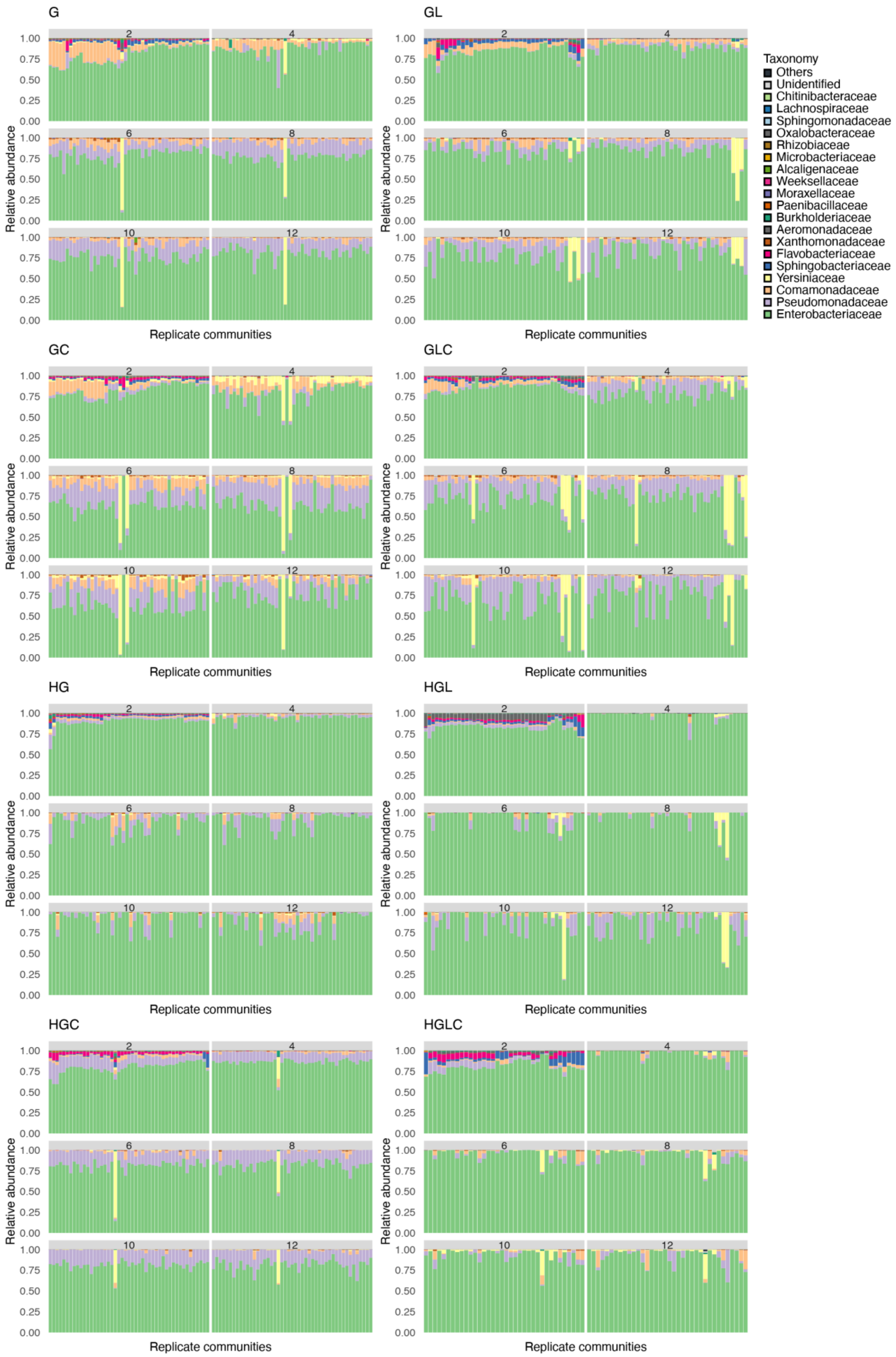
G
GL
GC
GLC
HG
HGL
HGC
HGLC
Relative abundance
Replicate communities
Taxonomy
Others
Unidentified
Chitinibacteraceae
Lachnospiraceae
Sphingomonadaceae
Oxalobacteraceae
Rhizobiaceae
Microbacteriaceae
Alcaligenaceae
Weeksellaceae
Moraxellaceae
Paenibacillaceae
Burkholderiaceae
Aeromonadaceae
Xanthomonadaceae
Flavobacteriaceae
Sphingobacteriaceae
Yersiniaceae
Comamonadaceae
Pseudomonadaceae
Enterobacteriaceae

51

**Fig. S6 | Variation in community structure among replicate samples (family level).** For each replicate community in each experimental treatment, changes in family-level community compositions (relative abundance) are shown. The numbers shown at the top of the bar plots refer to time points (days). The replicate samples were ordered based on unweighted pair group method with arithmetic mean (UPGMA) analyses performed on Day 2 for respective experimental treatments. The order of replicate communities on successive days is the same as that on Day 2.

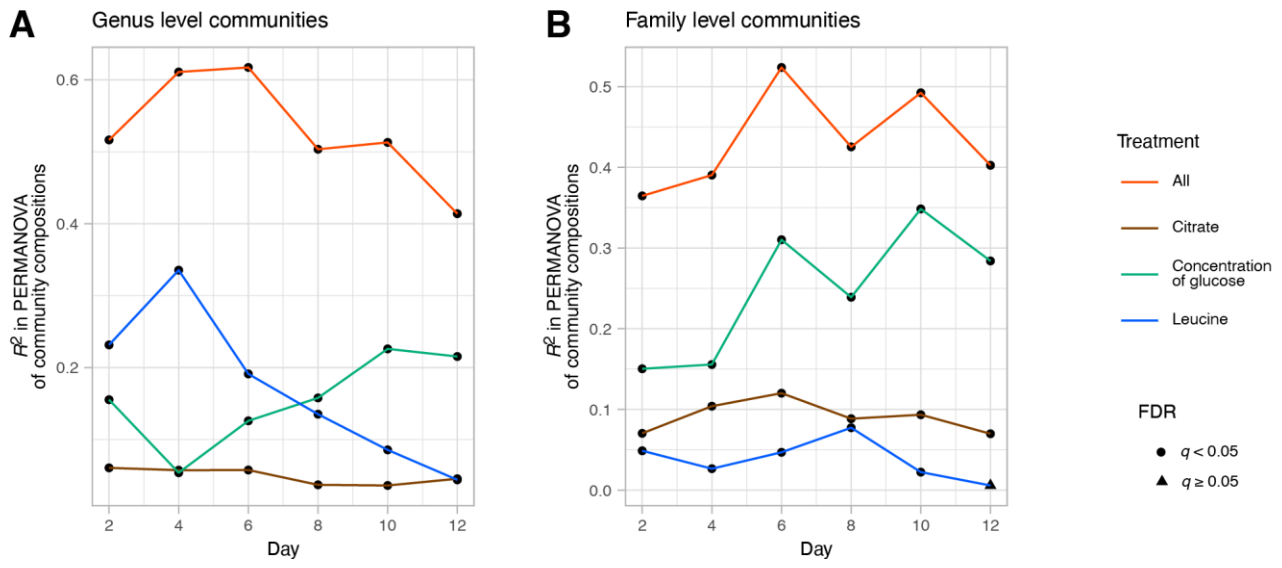


**Fig. S7 | Dependence of community structure on medium conditions (genus- and family-level analyses)**. **A** In each PERMANOVA model of genus-level community compositions, glucose concentration (high or low; df = 1), the presence/absence of leucine (df = 1), or the presence/absence of citrate (df = 1) was included as the explanatory variable. An additional model including all the medium conditions and interactions between them (df = 7) was examined as well. The coefficient of determination ($R^2$) is shown for each day. **B** PERMANOVA of family-level community compositions.

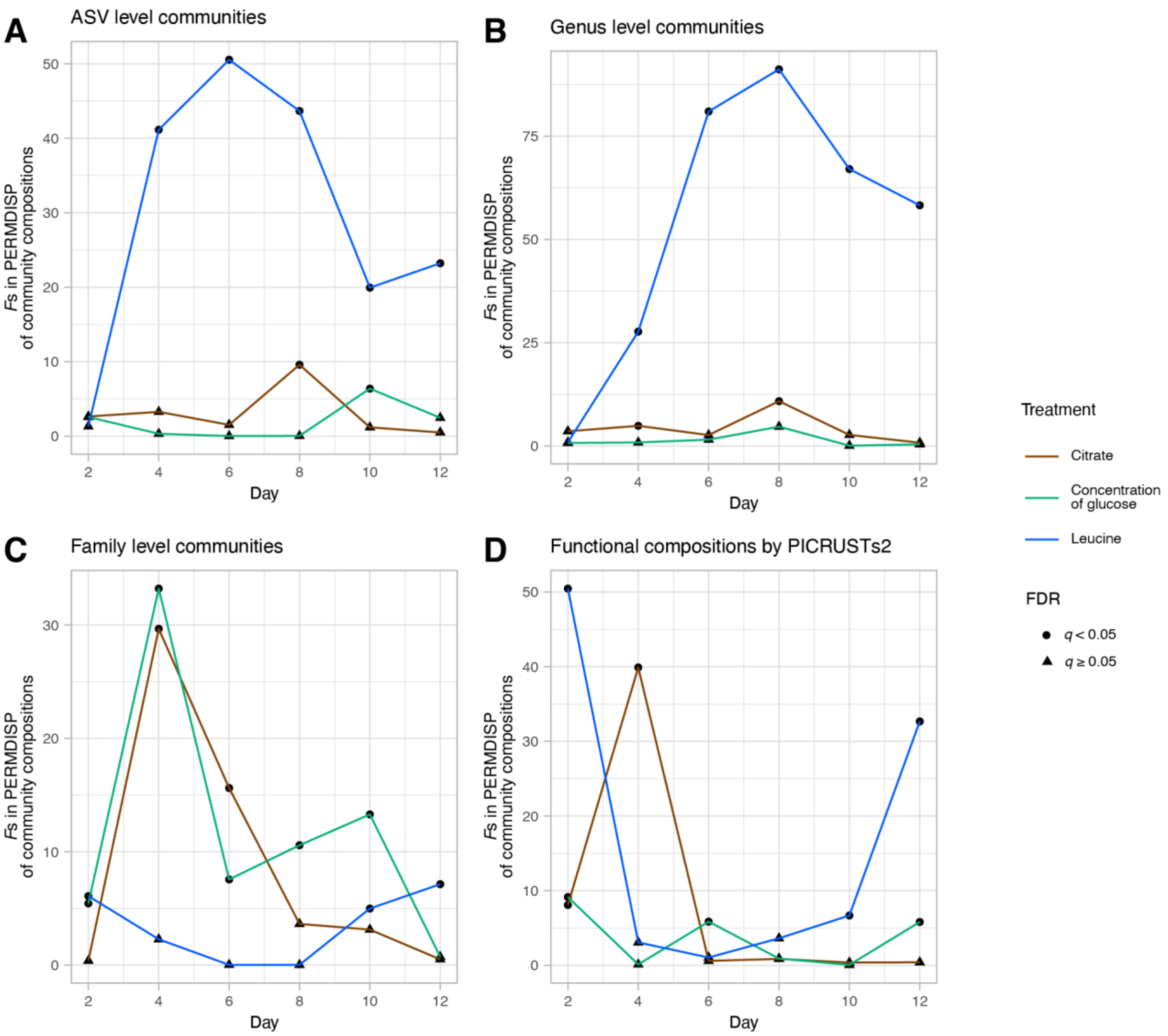


**Fig. S8 | Effects of medium conditions on community-structural dispersion among replicate samples. A** Potential effects of medium conditions on dispersion in community structure was examined by PERMDISP. In each PERMDISP model of dispersion of ASV compositions among replicate communities, glucose concentration (high or low; df = 1), the presence/absence of leucine (df = 1), or the presence/absence of citrate (df = 1) was included as the explanatory variable. The *F* statistics are shown for each day. **B** PERMDISP of genus-level compositions. **C** PERMDISP of family-level compositions. **D** PERMDISP of metabolic pathway compositions.

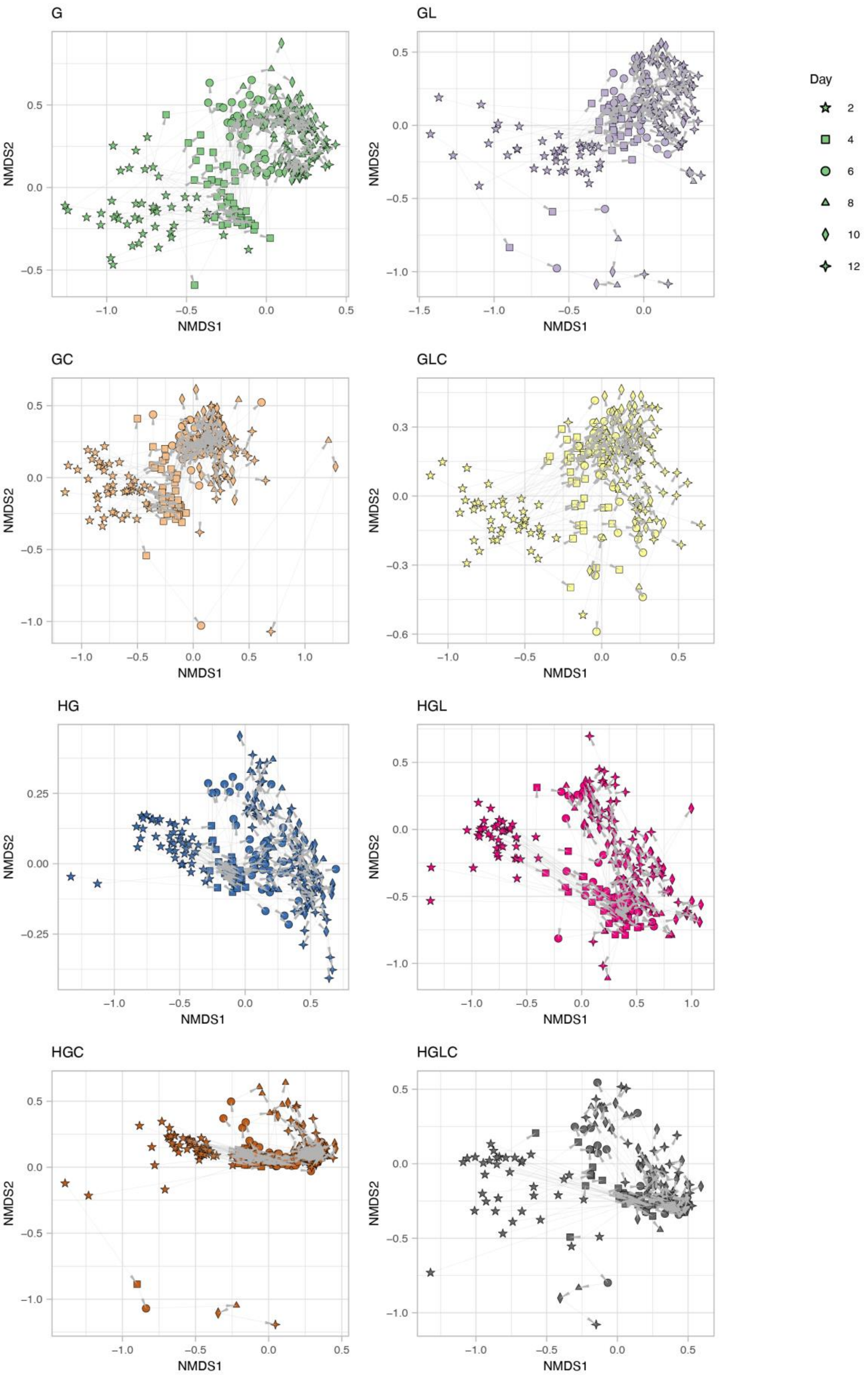
G
GL
GC
GLC
HG
HGL
HGC
HGLC
NMDS1
NMDS2
Day
2
4
6
8
10
12

**Fig. S9 | Time-series changes in community structure (genus level).** For each replicate community in each experimental treatment, time-series changes in family-level community structure are shown with arrows on the NMDS surface of the genus-level community compositions (stress = 0.165).

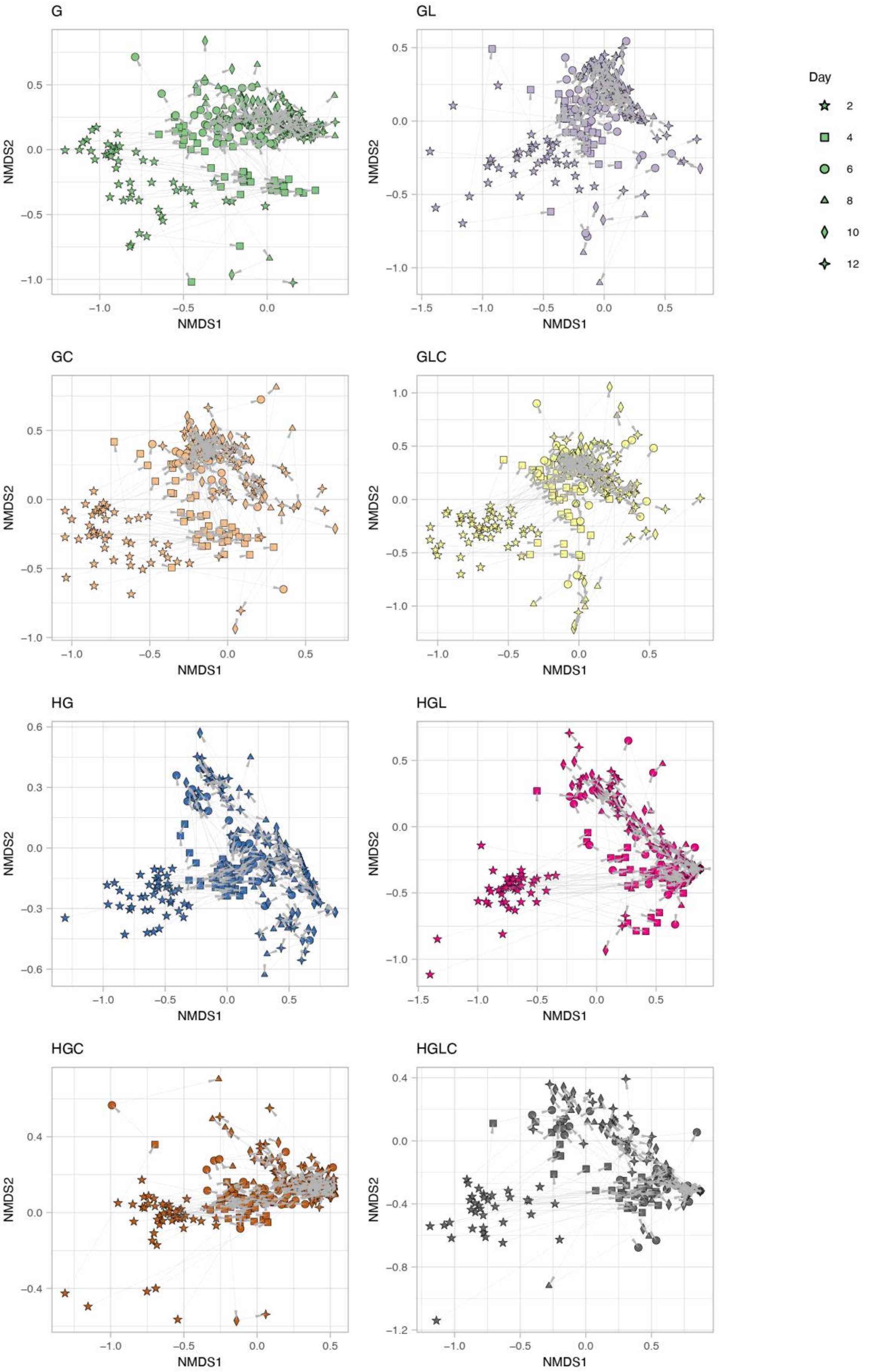
G
GL
GC
GLC
HG
HGL
HGC
HGLC
NMDS1
NMDS2
Day
2
4
6
8
10
12

**Fig. S10 | Time-series changes in community structure (family level).** For each replicate community in each experimental treatment, time-series changes in family-level community structure are shown with arrows on the NMDS surface of the family-level community compositions (stress = 0.148).

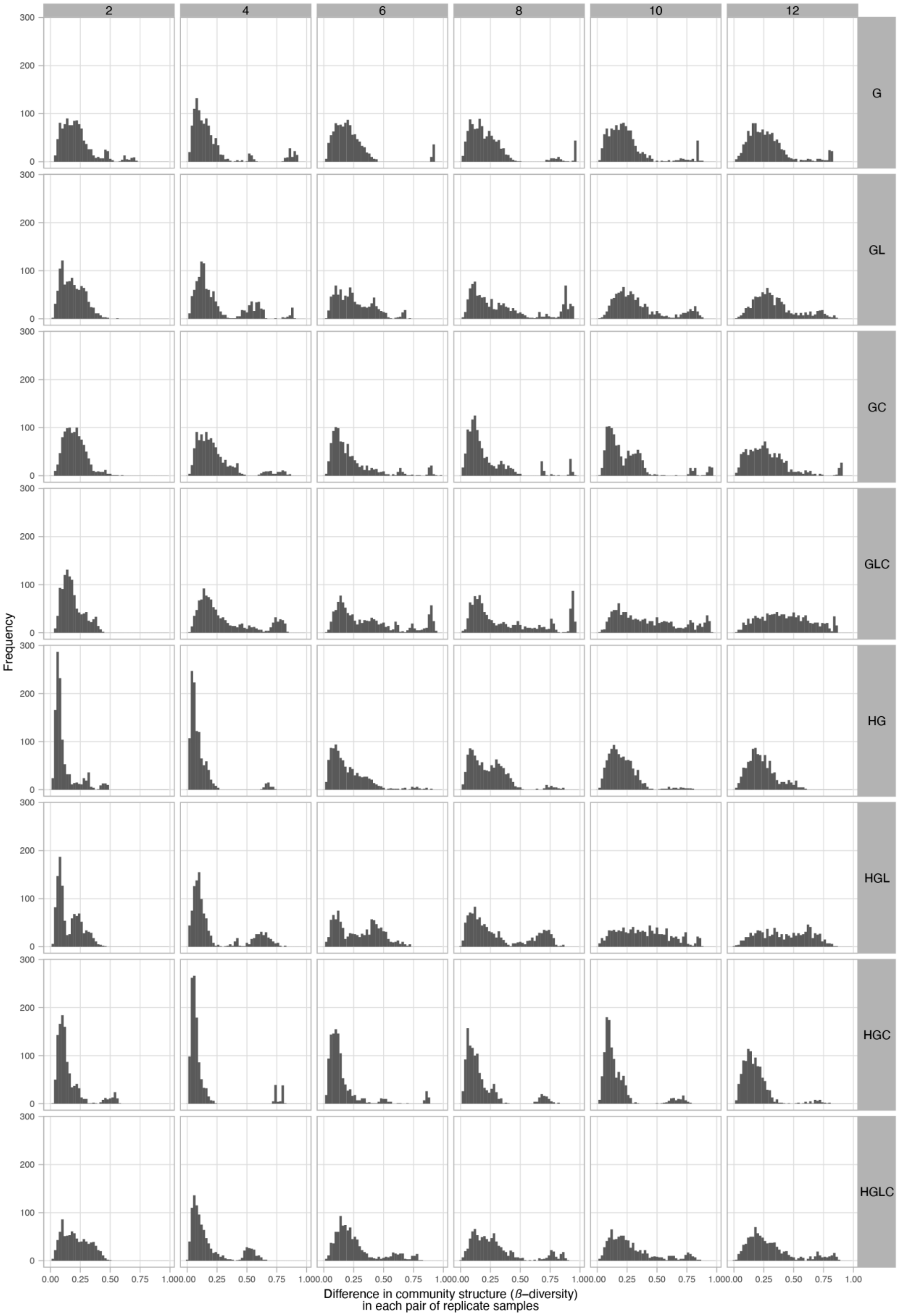
2
4
6
8
10
12
G
GL
GC
GLC
HG
HGL
HGC
HGLC
Frequency
0
100
200
300
0.00
0.25
0.50
0.75
1.00
Difference in community structure (β–diversity)
in each pair of replicate samples

78

**Fig. S11 | Histograms of community structural differentiation (genus level).** For each experimental treatment, difference in genus -level community structure (Bray-Curtis $\beta$-diversity) between replicate communities is shown as a histogram for each day. The numbers shown at the top of the histograms refer to the time points (days). The bi-modal or multi-modal distributions within these histograms suggest the presence of alternative community states.

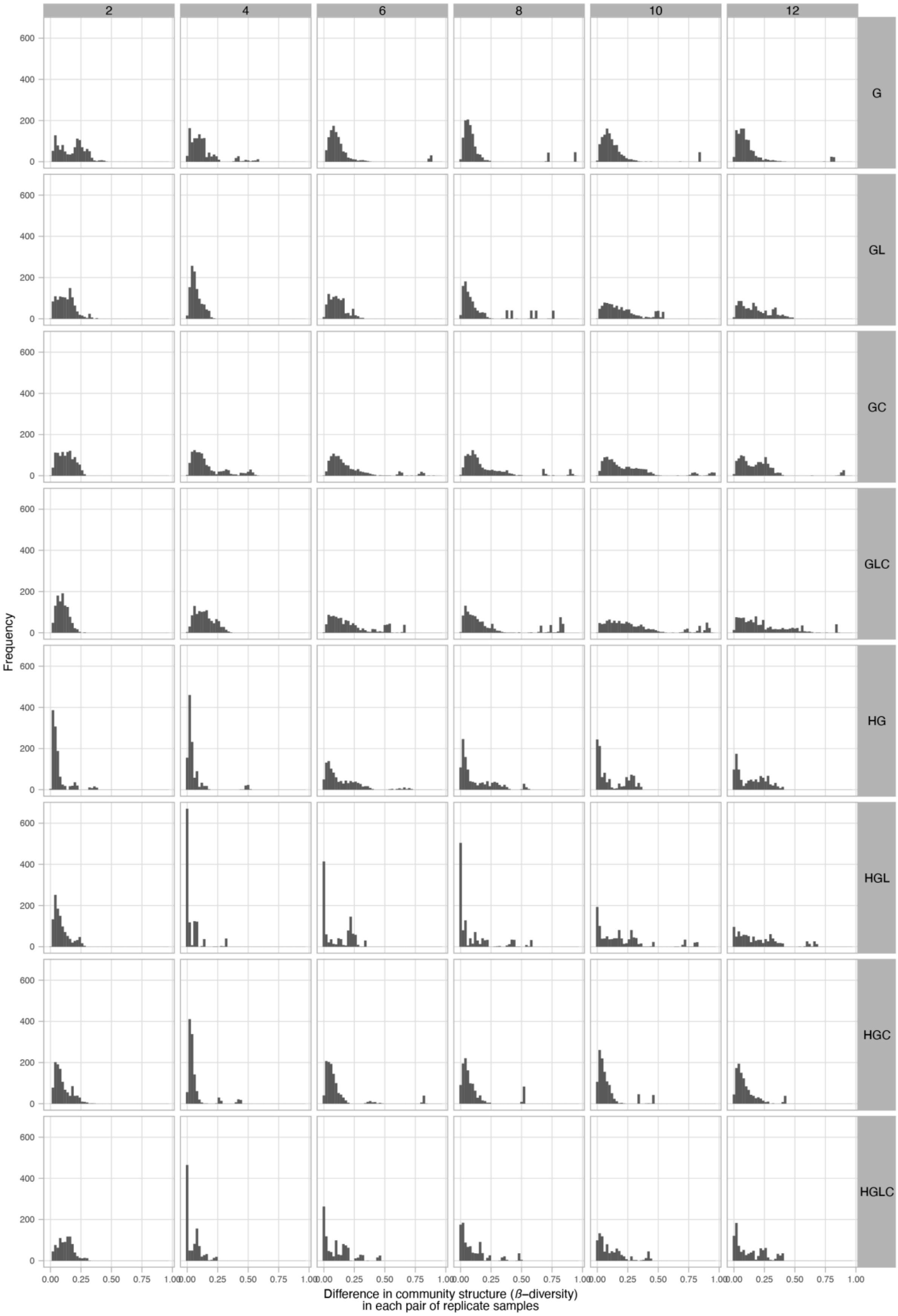

2
4
6
8
10
12
G
GL
GC
GLC
HG
HGL
HGC
HGLC
Frequency
0
200
400
600
0.00
0.25
0.50
0.75
1.00
Difference in community structure (β–diversity)
in each pair of replicate samples


83

**Fig. S12 | Histograms of community structural differentiation (family level).** For each experimental treatment, difference in family-level community structure (Bray-Curtis $\beta$-diversity) between replicate communities is shown as a histogram for each day. The numbers shown at the top of the histograms refer to the time points (days). The bi-modal or multi-modal distributions within these histograms suggest the presence of alternative community states.

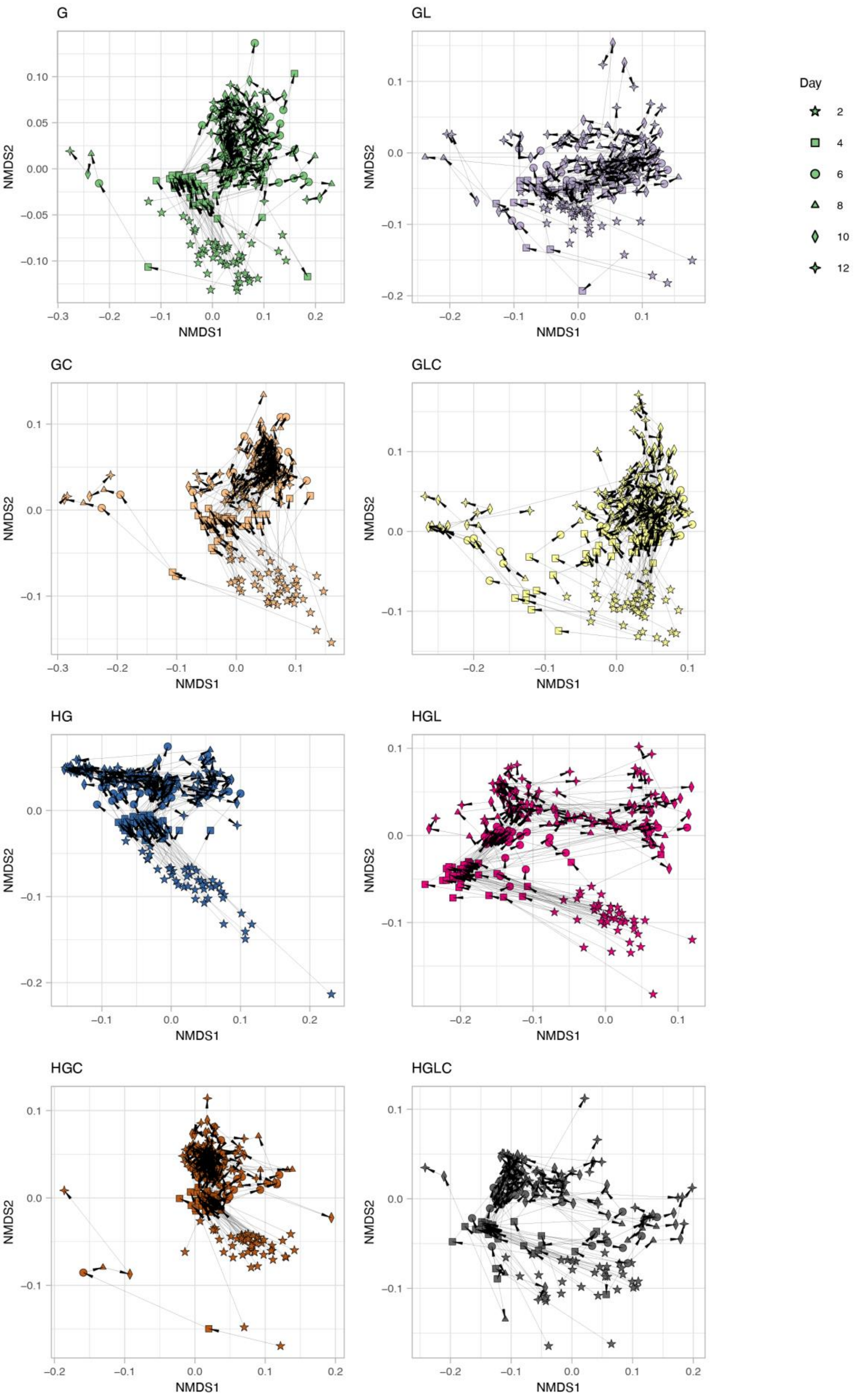


88

**Fig. S13 | Time-series changes in community functional profiles.** For each replicate community in each experimental treatment, time-series changes in community functional profiles (metabolic pathway/process compositions) are shown with arrows on the NMDS surface defined in Figure 7A (stress = 0.125).

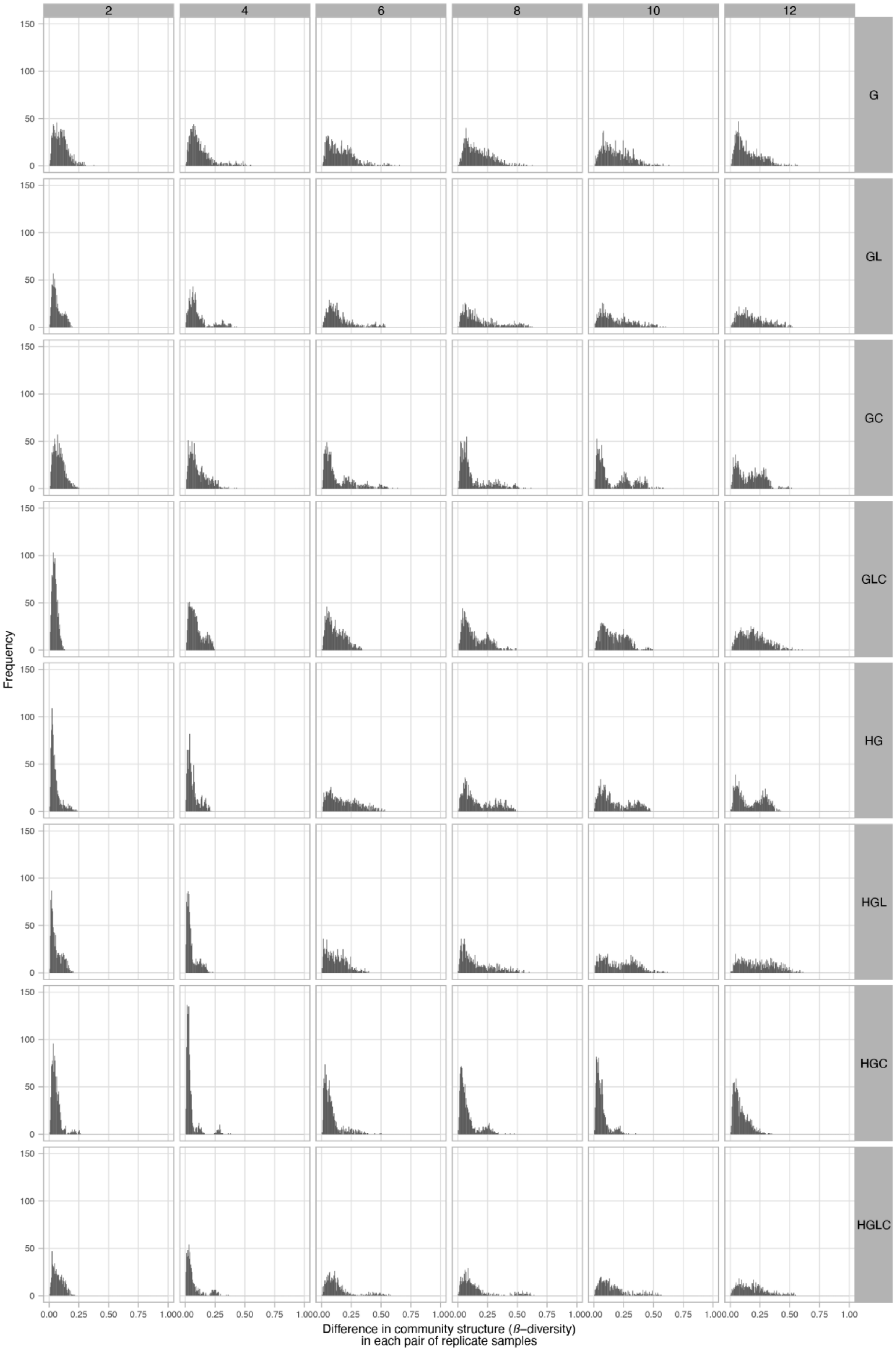
2
4
6
8
10
12
G
GL
GC
GLC
HG
HGL
HGC
HGLC
Frequency
150
100
50
0
0.00
0.25
0.50
0.75
1.00
Difference in community structure (β–diversity)
in each pair of replicate samples

92

**Fig. S14 | Histograms of differentiation in community functional profiles.** For each experimental treatment, difference in metabolic pathway/process compositions (Bray-Curtis $\beta$-diversity) between replicate communities is shown as a histogram for each day. The numbers shown at the top of the histograms refer to the time points (days). The bi-modal or multi-modal distributions within these histograms suggest the presence of alternative community states.

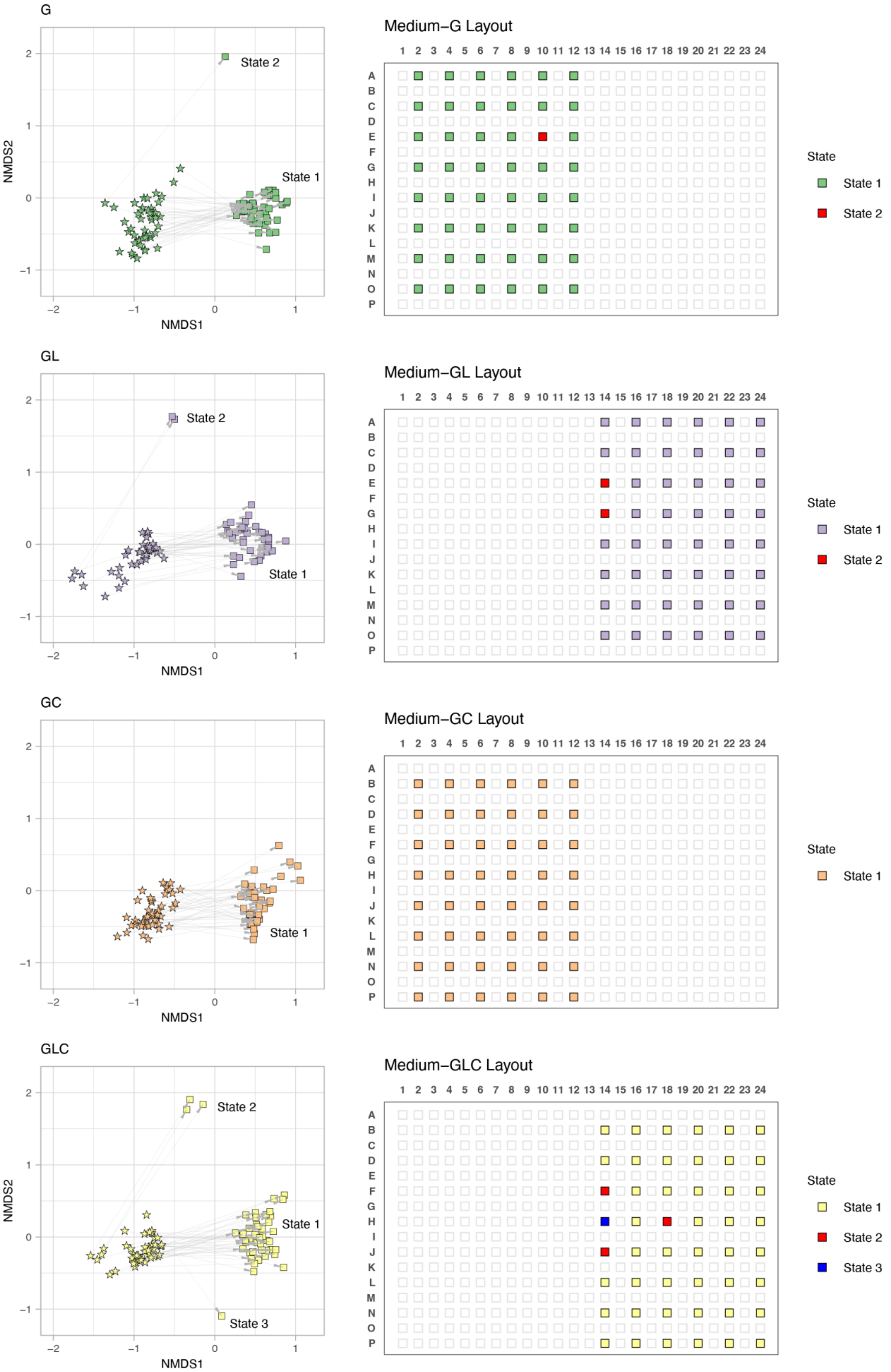

G
State 2
State 1
NMDS2
NMDS1
Medium−G Layout
State
State 1
State 2
GL
State 2
State 1
NMDS1
Medium−GL Layout
State
State 1
State 2
GC
State 1
NMDS1
Medium−GC Layout
State
State 1
GLC
State 2
State 1
State 3
NMDS2
NMDS1
Medium−GLC Layout
State
State 1
State 2
State 3

**Fig. S15 | Layout of replicate communities within the culture plate (Medium G, GL, GC, and GLC).** For each experimental treatment (medium condition), the positions of replicate samples on the deep-well plate are shown. For simplicity, ASV-level community compositions on Day 2 and Day 6 are plotted on the NMDS surface. Samples seemingly representing alternative community states are indicated within each treatment.

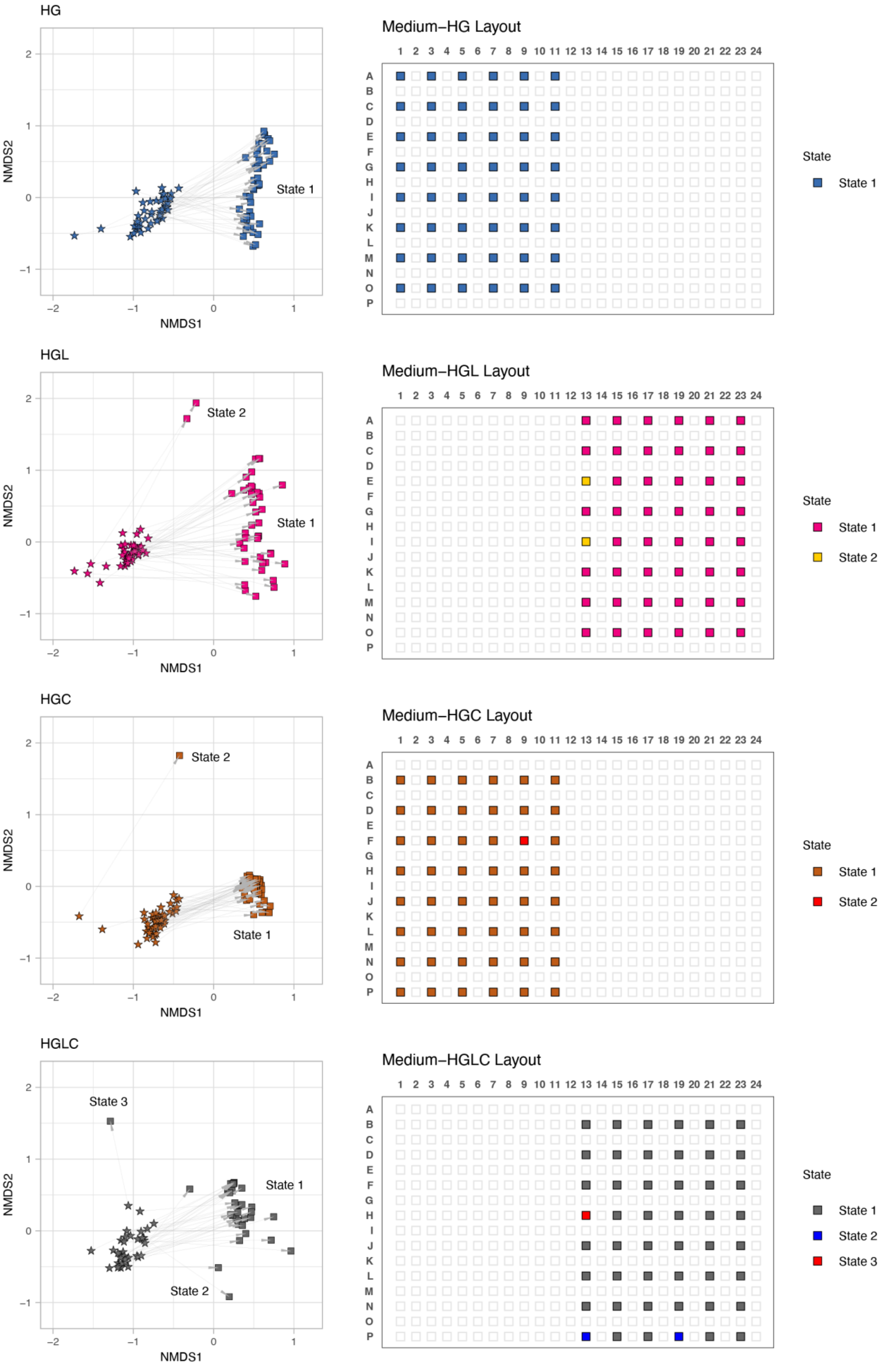
HG
State 1
NMDS1
NMDS2
Medium−HG Layout
State
State 1
HGL
State 2
State 1
NMDS1
NMDS2
Medium−HGL Layout
State
State 1
State 2
HGC
State 2
State 1
NMDS1
NMDS2
Medium−HGC Layout
State
State 1
State 2
HGLC
State 3
State 1
State 2
NMDS1
NMDS2
Medium−HGLC Layout
State
State 1
State 2
State 3

**Fig. S16 | Layout of replicate communities within the culture plate (Medium HG, HGL, HGC, and HGLC).** For each experimental treatment (medium condition), the positions of replicate samples on the deep-well plate are shown. For simplicity, ASV-level community compositions on Day 2 and Day 6 are plotted on the NMDS surface. Samples seemingly representing alternative community states are indicated within each treatment.
